# Estimating direct and indirect genetic effects on offspring phenotypes using genome-wide summary results data: a comparison of multivariate methods

**DOI:** 10.1101/2020.11.02.365981

**Authors:** Nicole M. Warrington, Liang-Dar Hwang, Michel Nivard, David M. Evans

## Abstract

Estimation of the direct genetic effect of an individuals’ own genotype on their phenotype, independent of any contaminating indirect parental genetic effects is becoming increasingly important. These conditional estimates are of interest in their own right, but are also useful for downstream analyses such as intergenerational Mendelian randomization. We compare several available multivariate methods that utilize summary results statistics from genome-wide association studies to determine how well they estimate conditional direct and indirect genetic effects, while accounting for sample overlap. Using robustly associated birth weight variants and data from the UK Biobank, we contrasted the point estimates and their standard errors at each of the individual loci compared to those obtained using individual level genotype data, in addition to comparing the computational time, inflation of the test statistics and number of genome-wide significant SNPs identified by each of the methods. We show that Genomic SEM outperforms the other methods in accurately estimating conditional genetic effects and their standard errors. We subsequently applied Genomic SEM to fertility data in the UK Biobank and partitioned the genetic effect into female and male fertility in addition to a sibling specific effect. This analysis identified one novel locus for fertility and replicated seven previously identified loci. We also identified genetic correlations between fertility and educational attainment, risk taking behaviour, autism and subjective well-being. We therefore recommend Genomic SEM to be used to partition genetic effects across the genome into direct and indirect components.

## Introduction

There is growing interest in estimating maternal genetic effects, via the intrauterine environment, on offspring outcomes (for example^1–3^) and also in elucidating the causal effect of maternal environmental exposures on offspring outcomes (for example,^3–5^). Likewise, it is also important to estimate the direct effect of an individual’s own genotype on their own phenotype independent of any indirect parental effects. These estimates can subsequently be used in Mendelian randomization studies to make causal inferences, without introducing biases due to dynastic effects and assortative mating^6–8^. However, for correct inference to be made regarding maternal genetic effects and an individual’s own genetic effect independent of parental effects, analyses must adjust for the individual’s own genetic effect on their own outcome as well as one or both parents. This adjustment has traditionally been performed using conditional analyses applied to individual level genotypes from mother-offspring pairs or parent-offspring trios; however, there are few cohorts worldwide with large numbers of genotyped mother-offspring pairs or parent-offspring trios with offspring phenotypes, leading to limited statistical power to detect an association in many studies. We have developed a structural equation model (SEM) that can partition genetic effects into maternal and offspring mediated components^9^. This model can incorporate data from genotyped mother-offspring pairs with offspring phenotypes, mother-offspring pairs with maternal genotypes and both mother and offspring phenotypes, individuals with their own genotype and phenotype and mothers with their own genotype and their offspring’s phenotype. We have recently described how the maternal and offspring partitioning from this SEM can be used to facilitate large-scale two-sample Mendelian randomization studies investigating whether maternal exposures are causally related to offspring outcomes^10^.

Although our SEM is flexible in terms of incorporating many study designs, it is computationally intensive when using individual level data, prohibiting its use for genome-wide association studies (GWAS). Therefore, we wanted to identify other existing methods that could be used on summary results statistics from GWAS to estimate the conditional maternal (paternal) and offspring genetic effects on a trait. There are a number of multivariate methods available that utilize summary statistics from GWAS of multiple traits. For example, metaCCA^11^, metaUSAT^12^, MTAG^13^, TATES^14^, S_HOM_ and S_HET_^15^, mtCOJO^16^ and most recently Genomic SEM^17^ are a subset of the multivariate methods that have been proposed for use with GWAS summary statistics to increase statistical power to detect an association with a correlated set of traits and diseases. Although we are interested in combining the summary statistics of the same trait from different genotypes (i.e. the individuals’ own genotype and their mother’s genotype), we hypothesize that some of these methods could be appropriate. However, because we are interested in using the adjusted maternal and offspring genetic effect in downstream analyses, such as Mendelian randomization, we need methods that would provide effect estimates (and standard errors) for each genetic variant. In addition, if we are to use publically available summary statistics from large GWAS, such a method would need to account for any known or unknown overlap of individuals contributing to maternal (paternal) and offspring GWAS.

In this manuscript, we compare several different multivariate methods to identify the most appropriate method for partitioning the genetic effect of a trait into maternal and offspring components, based on how well the effect estimates compare to those from our SEM using individual level data, the computational time and how well the method accounts for unknown sample overlap. We use birth weight to compare the different methods as we have a substantial number of known associated genetic loci for birth weight, with the genetic effect partitioned into maternal and offspring genetic components. We subsequently use the most appropriate method to conduct conditional GWAS of fertility, partitioning the effects into parental and offspring mediated components providing evidence for how these different loci exert their effect on number of children in a family.

## Methods

### Selecting statistical methods

We searched the literature for multivariate methods that fit the following four criteria: 1) had published code or software, 2) used summary results statistics and did not require individual level data, 3) accounted for sample overlap and 4) produced an effect size estimate and standard error for each trait. In addition to our published structural equation model (SEM)^9^ and linear approximation of the SEM^3^, we identified three published methods including multi-trait analysis of GWAS (MTAG)^13^, multi-trait-based conditional and joint analysis using GWAS summary data (mtCOJO)^16^ and Genomic SEM^17^. Although MTAG and mtCOJO are not specifically designed to partition genetic effects into maternal and offspring components, we were interested in investigating whether they would approximate the effects accurately.

### Participants

The UK Biobank has ethical approval from the North West Multi-Centre Research Ethics Committee (MREC), which covers the UK, and all participants provided written informed consent. UK Biobank phenotype data was available on 502,543 individuals, of which 280,142 reported their own birth weight at either the baseline or first two follow-up visits. There were 7,701 individuals who were part of a multiple birth and were excluded from the analyses. There were 10,670 individuals who reported their birth weight at more than one visit, with 83 individuals reporting the two values to be different by more than 1kg; these individuals were excluded from the analyses. For those individuals who reported different values between baseline and follow-up (<1kg) we took the measure from the first reported visit for the analyses. Finally, we excluded individuals who reported their birth weight to be <2.5kg or >4.5kg, as these are implausible for live term births before 1970. In total, 234,154 individuals had data on their birth weight matching our inclusion criteria.

Women in the UK Biobank were also asked to report the birth weight of their first child to the nearest pound and were converted to kilograms for analyses (N=216,782). We excluded individuals with multiple measures that differed by >1kg (N=29) or if their birth weight was <2.2kg (5 pounds) or >4.6kg (10 pounds), leaving 210,423 individuals with birth weight of their first child matching our inclusion criteria.

We analysed genetic data from the April 2018 release of imputed data from the UK Biobank, a resource that is described extensively elsewhere^18^. In addition to the quality control metrics performed centrally by the UK Biobank, we defined a subset of “white European” ancestry, unrelated individuals. First, we generated ancestry informative principal components (PCs) in the 1000 genomes samples. The UK Biobank samples were then projected into this PC space using the single nucleotide polymorphism (SNP) loadings obtained from the PC analysis using the 1000 genomes samples. The UK Biobank participants’ ancestry was classified using K-means clustering centered on the three main 1000 genomes populations (European, African, South Asian). Those clustering with the European cluster were classified as having European ancestry. The UK Biobank participants were asked to report their ethnic background. Only those reporting as either “British”, “Irish”, “White” or “Any other white background” were included in the clustering analysis. Secondly, to identify a subset of unrelated individuals in the UK Biobank, we generated a genetic relationship matrix in the GCTA software package^19^ (version 1.90.2) and excluded one of every pair of related individuals with a genetic relationship greater than 9.375%. A subset of 257,696 individuals with genotype data, a valid birth weight for themselves or their first child, were unrelated and were genetically of ‘white European’ ancestry remained for analysis. Of these, 72,274 were men so only reported their own birth weight, 25,951 women reported only their own birth weight, 73,968 reported only the birth weight of their first child and 85,503 reported both. We adjusted both the individuals’ own birth weight and the birth weight of their first child for the principal components provided by the UK Biobank, assessment center and genotyping array, and sex for own birth weight, and then created z-scores.

Because we were interested in how each of the statistical methods handled sample overlap, we created two sets of data (**Figure 1**). The first contained all of the data available, including the 85,503 that contributed to both the GWAS of own birth weight (N=183,728) and the GWAS of offspring birth weight (N=159,471). The second contained all of the data for the GWAS of offspring birth weight (N=159,471) but excluded those individuals from the GWAS of own birth weight that were included in the GWAS of offspring birth weight (N=72,274 + 25,951 = 98,225).

**Figure 1:**
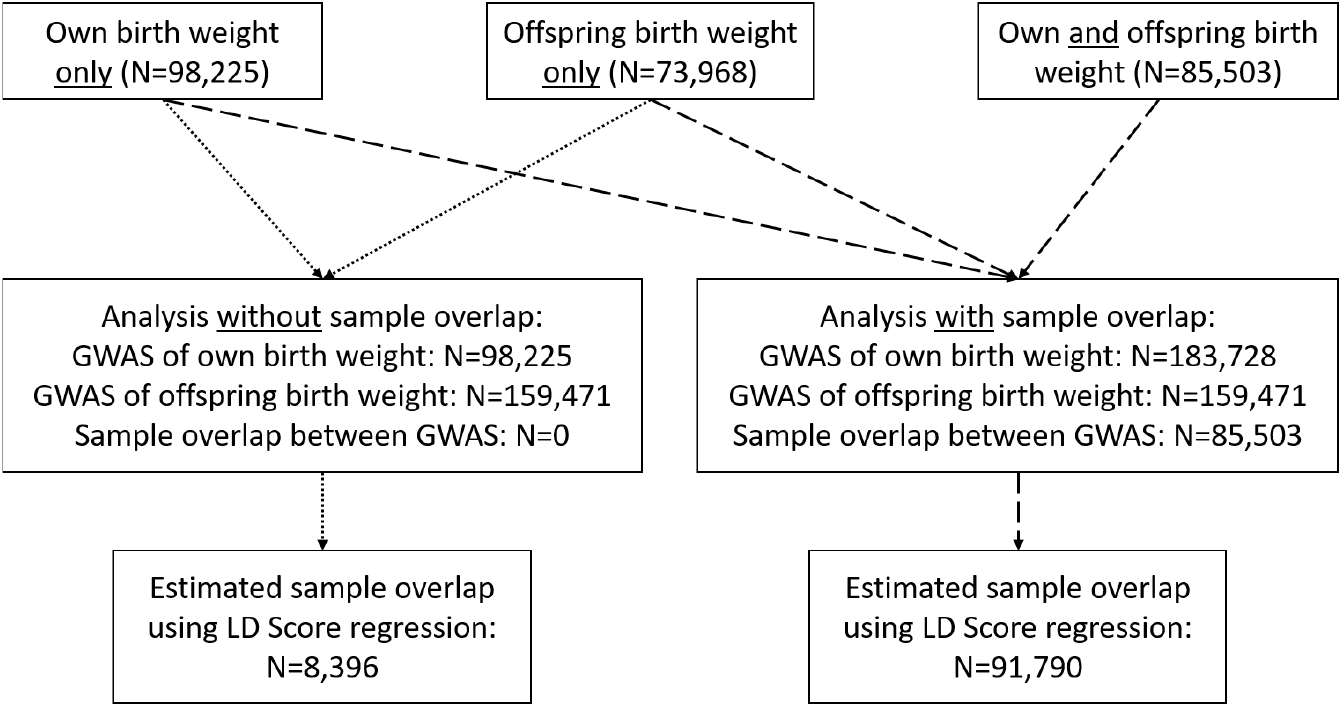
Schematic of the study design for comparing methods using self-reported birth weight data from the UK Biobank. We conducted two sets of analysis, one with and one without sample overlap between the genome-wide association studies (GWAS), to investigate the effect of sample overlap in each of the methods.

### GWAS analysis

GWAS of own and offspring birth weight was conducted using a linear mixed model implemented in BOLT-LMM v2.3.2^20^ to account for population structure and subtle relatedness. Only autosomal genetic variants which were common (minor allele frequency > 0.1%), had Hardy-Weinberg equilibrium P-value>1×10^-6^ and missingness < 0.1 were included in the genetic relationship matrix (GRM). We excluded genetic variants with an INFO score < 0.4 and minor allele frequency < 0.1% from the analysis in BOLT-LMM (BOLT-LMM uses the full sample to exclude SNPs based on these thresholds, so some SNPs may have minor allele frequency <0.1% in our subset of the UK Biobank data with clean birth weight data). We then used the summary statistics from the GWAS of own and offspring birth weight in the following analyses to estimate the conditional maternal and offspring genetic effects at each genetic variant.

### SEM analysis using summary statistics

The SEM to estimate the adjusted maternal and offspring genetic effects has been described in detail previously^9^ (**Supplementary Figure 1**). Briefly, to estimate the parameters for the adjusted offspring and maternal genetic effects on birth weight, we use three observed variables available in the UK Biobank; the participant’s genotype, their own self-reported birth weight, and in the case of the UK Biobank women, the birth weight of their first child. Additionally, the model comprises two latent (unobserved) variables, one for the genotype of the UK Biobank participant’s mother and one for the genotype of the participant’s offspring. From biometrical genetics theory, these latent genetic variables are correlated 0.5 with the participant’s own genotype, so we fix the path coefficients between the latent and observed genotypes to be 0.5. We have previously described how the SEM can be fit with either the individual level data or observed covariance matrices derived from the individual level data^9^. To derive the observed covariance matrices, we need the allele frequency of the genetic variant, beta coefficient from the regression model of the genetic variant on own or offspring phenotype, variance of the phenotype (which will be one if the phenotype was standardized prior to the regression analysis) and the sample size. We assume that the allele frequency, beta coefficient and variance is consistent across the following three groups, but the sample size will differ: individuals with their own phenotype only, individuals with their offspring’s phenotype only and individuals with both. We therefore need to estimate the sample overlap from the summary statistics in order to include the sample size for each of the covariance matrices. To do this, we performed bivariate linkage disequilibrium (LD) score regression (version 1.0.0) analysis using the summary statistics from the GWAS of own birth weight and the GWAS of offspring birth weight and used the regression intercept to estimate the number of individuals in both analyses. We used a phenotypic correlation between own and offspring birth weight of 0.24 in the calculation, which was estimated using the cleaned phenotype data that was included in the GWAS analysis. We have previously shown that the SEM has difficulty optimizing with low frequency variants, so we excluded SNPs with a minor allele frequency less than 0.5%. For each genetic variant we then calculated the observed covariance matrices from the summary statistics and fit the SEM with the relevant estimated sample sizes. We calculated a Wald P-value for the maternal and offspring genetic effects using the effect size estimates and their standard errors. We conducted the analysis of each chromosome in parallel to reduce the computational time. Analyses were conducted in R (version 3.4.3) using the OpenMx package (version 2.6.9).

### Analysis using a linear approximation of the SEM

We have previously derived a weighted linear model that is a good approximation of the SEM but substantially less computationally intensive^3^. This model uses a linear transformation of the effect sizes from the GWAS of own birth weight and the GWAS of offspring birth weight based on the principles of ordinary least squares linear regression. The offspring effect at each genetic variant is estimated as:

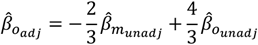

And the corresponding standard error is:

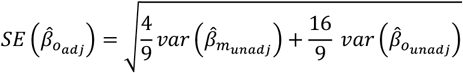

Where 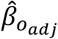 is the offspring effect adjusted for the maternal effect, 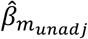 is the unadjusted maternal effect from the GWAS of offspring birth weight and 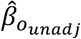 is the unadjusted offspring effect from the GWAS of own birth weight. Likewise, the maternal effect is estimated as:

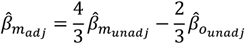

And standard error is:

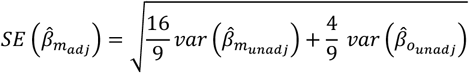

Where 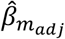 is the maternal effect adjusted for the offspring effect. The full derivation can be found in Warrington et al. (2019)^3^. Similar to the SEM using summary statistics, we calculated a Wald P-value for the maternal and offspring genetic effects using the effect size estimates and their standard errors. This method assumes that the two unadjusted GWAS are independent, and do not contain any sample overlap. We performed the linear transformation in R (version 3.4.3) and each of the chromosomes were run in parallel to reduce the computational time.

### Analysis using MTAG

MTAG enables joint analysis of multiple traits using summary statistics from GWAS, while accounting for the possibility of overlapping samples, and produces trait-specific effect estimates for each genetic variant^13^. It is based on the idea that when GWAS estimates from different traits are correlated, the SNP effect estimates can be improved by incorporating information from the other correlated traits. This method was not developed to estimate SNP effects conditional on parental genotypes; however, we were interested in investigating how well it approximated the maternal and offspring genetic effects on birth weight. We used python version 2.7.12 and set the lower sample size bound to zero (--n_min 0.0) when running MTAG.

### Analysis using mtCOJO

mtCOJO^16^ performs an approximate multi-trait-based conditional GWAS using summary statistics from a GWAS of two or more traits. For our birth weight analysis, the method approximates the following two models:

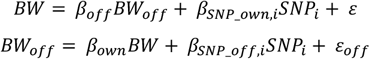

Where *BW* is the individual’s own birth weight, *BW_off_* is offspring birth weight, *SNP_i_* is the i^th^ SNP, *β_off_* is the effect of offspring birth weight on the an individual’s own birth weight, *β_SNP_own,I_* is the effect of SNP_i_ on the individual’s own birth weight, *β_own_* is the effect of individual’s own birth weight on the their offspring’s birth weight, *β_SNP_off,I_* is the effect of SNP_i_ on offspring birth weight, ε and ε_off_ are the residuals. Although this method conditions on the phenotype of the other individual in the pair (i.e. conditioning on the offspring’s phenotype when analysing own birth weight, rather than their genotype), we wanted to investigate how well this approach would approximate conditioning on the genotype. In other words, we were interested in investigating how well 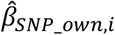 approximates 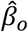 from the SEM for SNP *i*, and 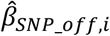 approximates 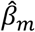.

To conduct analysis in mtCOJO, we needed a reference sample with individual level genotypes for LD estimation. Therefore, we randomly sampled 50,000 individuals from the UK Biobank and extracted their imputed genetic data, using Plink2 (released 18 March 2019), for each of the unique genetic variants that were included in the cleaned GWAS summary statistics. mtCOJO could not handle genetic variants that had the same rs number but different alleles (i.e. multi-allelic markers), so we removed all duplicate rs numbers from the reference dataset. Using this reference dataset, we conducted the mtCOJO analysis with the default parameters in GCTA (version 1.92.0beta3), using the summary statistics from the GWAS of own birth weight as the outcome and conditioning on offspring birth weight and then using the summary statistics from the GWAS of offspring birth weight as the outcome and conditioning on own birth weight.

### Analysis using Genomic SEM

Genomic SEM^17^ is a multivariate method for analysing the joint genetic architecture of traits using GWAS summary statistics. It is a very flexible method based on structural equation modelling of a genetic covariance matrix derived from summary results GWAS data and can be used to estimate conditional maternal (paternal) and offspring specific genetic effects. The specific model that we fit for the birth weight data is depicted in **Supplementary Figure 2**.

We conducted GWAS analysis using the userGWAS function in Genomic SEM v0.0.2 (installed 9^th^ Jan 2020), which creates genetic covariance matrices for individual SNPs and estimates SNP effects for a user specified multivariate GWAS. Following the workflow described on the github Wiki, we ran multivariate LD score regression to estimate the genetic covariance matrix and corresponding sampling covariance matrix, which accounts for any potential sample overlap between the GWAS summary statistics. After preparing the summary statistics for analysis, we used the estimated matrices to run the GWAS using 50 cores on a computing cluster.

### Comparison of methods

We fit the same SEM as described above, but using the individual level data rather than observed covariance matrices, for the 300 autosomal genetic variants that reached P<5×10^-8^ in the latest GWAS of birth weight^3^; we excluded rs2428362 from the comparison as it is tri-allelic. For the SEM using the data with no sample overlap (i.e. where the individuals from the UK Biobank had either their own birth weight measure or their offspring’s, but not both), we did not estimate the correlation between the birth weight measures as we had no data to estimate the parameter. We visualized the difference between the effect size estimates and standard errors from this SEM and those estimated using the GWAS summary statistics and methods described above. We conducted a heterogeneity test to assess the difference in the beta coefficients using the rmeta package (version 3.0) in R (version 3.5.2).

We were also interested in how the methods compared in terms of computational time, inflation of the test statistics and number of genome-wide significant SNPs identified. We saved the computational time for each of the methods (we used the run time from chromosome two for the time of the SEM using summary statistics as each chromosome ran in parallel so this was the longest chromosome to run) for comparison purposes. We note that the computational time will differ between computing resources and we present them here to compare the methods relative to each other. We conducted an LD score regression (version 1.0.0) analysis to estimate the inflation in test statistics for each method.

### Simulations under the null

In the methods comparison, it appeared that mtCOJO had increased statistical power to detect both the maternal and offspring specific genetic effects. Given this comparison used SNPs that were known to be associated with birth weight, we were interested if the method had increased type one error rate under the null hypothesis of no genetic effect. We simulated own and offspring birth weight based on 294 SNPs on chromosomes 1-21 that reached P<5×10^-8^ in the latest GWAS of birth weight^3^ and analysed the data from chromosome 22 (i.e. where there were no SNPs associated with the simulated birth weight data). Assuming autosomal Mendelian inheritance, additivity and unit variance, latent variables for the genotype of the individual’s mother (i.e., grand-maternal genotype) and the individual’s offspring’s genotype were generated, based on the individual’s own genotype at each of the 294 SNPs. The individual’s own birth weight variable (BW) for each family *i*, was generated using the following equation:

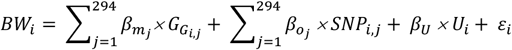

where *β_m_j__* denotes the maternal genetic effect at SNP *j, G_G_i,j__* is a latent variable indexing the genotype of individual *i*’s mother at SNP *j, β_O_j__* denotes the offspring genetic effect at SNP *j, SNP_i,j_* is the genotype of the *i*th individual at the *j*th SNP, *U* is a standard normal random variable representing all residual genetic and environmental sources of similarity between mother and offspring, *β_U_* is the total effect of *U* on the individual’s own birth weight and was fixed to 0.5, and *ε* is a random normal variable with mean zero and variance needed to ensure that BW has unit variance asymptotically. Similarly, offspring birth weight (BW_O_) for each family *i*, was generated using the following equation:

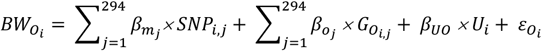

where *G_O_i,j__* is a latent variable indexing the offspring genotype, *β_UO_* is the total effect of *U* on offspring birth weight and was fixed to 0.5, and *ε_O_* is a random normal variable with mean zero and variance needed to ensure that BW_O_ has unit variance asymptotically. For *β_m_j__* and *β_o_j__* at each SNP we used the maternal and offspring specific effects estimated using a structural equation model in the recent GWAS of birth weight^3^; we added 0.01 to each effect size to ensure the effect sizes were large enough to detect in UK Biobank alone.

We adjusted both the simulated values of own birth weight and offspring birth weight for the principal components provided by the UK Biobank and genotyping array, then created z-scores. To closely mimic the real data analysis, we set the simulated birth weight values to missing if the participants in UK Biobank did not have birth weight or offspring birth weight available for analysis. We generate the same two sets of data; the first included all individual who had birth weight and offspring birth weight (which included sample overlap) and the second included all individuals with offspring birth weight but excluded those individuals from the GWAS of own birth weight that were included in the GWAS of offspring birth weight (i.e. no individuals were in both analyses). We conducted a GWAS in BOLT-LMM v2.3.2^20^ for both the simulated own and offspring birth weight, using the same GRM as was used in the birth weight analyses, and excluding genetic variants with an INFO score < 0.4 and minor allele frequency < 0.1%. As we were specifically interested in the performance of mtCOJO when there are no significant loci, we subsequently conducted analysis of the simulated data from chromosome 22 in mtCOJO using the same methods as the main analysis. For comparison, we also conducted analysis using the linear approximation of the SEM.

### Application to “real” data: Fertility GWAS

In the UK Biobank, 272,579 women and 225,349 men reported how many children they had given birth to (live births only) or fathered respectively. Additionally, each of the participants reported how many full brothers (N= 493,181) and sisters (N= 493,257) they had at each follow-up. There were 28,609 women who reported how many children they mothered at more than one visit, 263 (0.9%) of whom their response changed over time so were excluded. Similarly, there were 26,171 men who reported how many children they had fathered at more than one visit, 1,103 (4%) of whom their response changed over time so were excluded. In terms of siblings, 54,480 participants reported how many full brothers they had and 54,489 how many full sisters they had at more than one visit, with 2,430 (4%) and 1,802 (3%) excluded respectively because their response changed over time. We added the number of brothers and sisters to get the total number of siblings, with 489,701 participants reporting how many siblings they had available for analysis. Participants reporting greater than 10 siblings (N=1,720, 0.4%) or children (mothered N=18, 0.007%; fathered N=43, 0.02%) were recoded to have 10 in case these were data errors and to avoid a distribution with a large tail. We excluded individuals who were not part of our ‘white European’ ancestry cluster, leaving 237,768 women reporting how many children they mothered, 199,570 men reporting how many children they had fathered and 430,466 individuals reporting how many siblings they have available for GWAS analysis.

GWAS of the number of siblings and number of children for the women and men were conducted using a linear mixed model implemented in BOLT-LMM v2.3.2^20^. We used the same GRM as was used in the birth weight analyses, and excluded genetic variants with an INFO score < 0.4 and minor allele frequency < 0.1% from the analysis. We adjusted for the 40 principal components provided by the UK Biobank, genotyping array, age that the number of siblings or children was reported, assessment centre that the participant attended, and for the siblings analysis we also adjusted for sex of the participant. Subsequently, we used the summary statistics to conduct two analyses in Genomic SEM;

1. Similar to the birth weight analysis, we used the GWAS summary statistics of the number of siblings and number of children mothered to generate female and sibling specific genetic effects on fertility (**Supplementary Figure 3A**).
2. We used the GWAS summary statistics for all three traits to generate female, male and sibling specific effects for fertility to illustrate how the structural equation model can be extended when data from fathers is also available (**Supplementary Figure 3B**).

To investigate how Genomic SEM performed on a genome-wide scale, we estimated genetic correlations between the unconditional GWAS conducted in BOLT-LMM and the conditional GWAS conducted in Genomic SEM using LD score regression^21^. Subsequently, we used LD Hub^22^ (ldsc.broadinstitute.org) to estimate genetic correlations between the conditional estimates from Genomic SEM and a range of developmental, reproductive, behavioural, neuropsychiatric and anthropometric phenotypes that were investigated in Barban *et al*^23^. We also investigated the genetic correlation between the conditional estimates and risk taking behaviour as one of the genome-wide significant loci was previously associated with this trait. Due to the different linkage disequilibrium structure across ancestry groups, we only used summary statistics from LD Hub that were from European origin. There were several traits that had summary statistics from multiple GWAS available in LD Hub, so we used the latest GWAS to estimate the genetic correlations with fertility.

## Results

Approximately 19 million genetic variants were included in the GWAS analysis of own and offspring birth weight that passed our filtering criteria (INFO score > 0.4 and minor allele frequency < 0.1%). We excluded variants from the SEM using summary statistics if the minor allele frequency in the sample was < 0.5%; this lead to approximately 11 million genetic variants with results. MTAG also implements additional filtering criteria; variants with missing values, variants that are not SNPs, variants with duplicated rs numbers and variants that are strand ambiguous are excluded leading to approximately 14 million genetic variants with results.

We used bivariate LD score regression^21^ to estimate the sample overlap between the GWAS of own and offspring birth weight. We observed a regression intercept of 0.1287 (0.0078) in the analyses where there were individuals in both the GWAS of own and offspring birth weight, indicating that approximately 91,790 individuals were in both GWAS (true overlap is 85,503 individuals). In the analyses where there were unique individuals in the GWAS of own and offspring birth weight, the observed regression intercept from LD score regression was 0.0161 (0.0064), indicating that approximately 8,396 individuals were in both GWAS (true overlap is 0 individuals). These estimates of sample overlap were used in the SEM analysis using covariance matrices derived from the GWAS summary statistics.

### Comparison with SEM using individual level data

We compared the effect size and standard errors estimated using the SEM with the individual level data to those estimated using methods based on the summary statistics from the GWAS of own and offspring birth weight for the 300 autosomal genome-wide significant SNPs identified in the latest GWAS of birth weight^3^. Due to the additional exclusions, MTAG had 258 SNPs available for comparison and the SEM using summary statistics had 298 SNPs. We use the estimates from the SEM using individual level data as a baseline comparator as we have previously shown that they are asymptotically unbiased estimates of the maternal and offspring genetic effects^9^. **Figure 2** indicates that the effect sizes for each of the 300 genetic variants are accurately estimated using the SEM based on covariance matrices derived from the summary statistics, the linear approximation of the SEM or Genomic SEM, and these do not appear to be influenced by sample overlap. Given mtCOJO and MTAG were not developed to estimate maternal and offspring specific genetic effects, it is perhaps not surprising that there appears to be a slight underestimation of the effect sizes using mtCOJO, which is consistent with and without sample overlap. In contrast, the effect size estimates from MTAG (which also is not explicitly developed for estimating maternal and offspring genetic effects) differ from the SEM effect sizes for both the maternal and offspring effect, with and without sample overlap.

**Figure 2:**
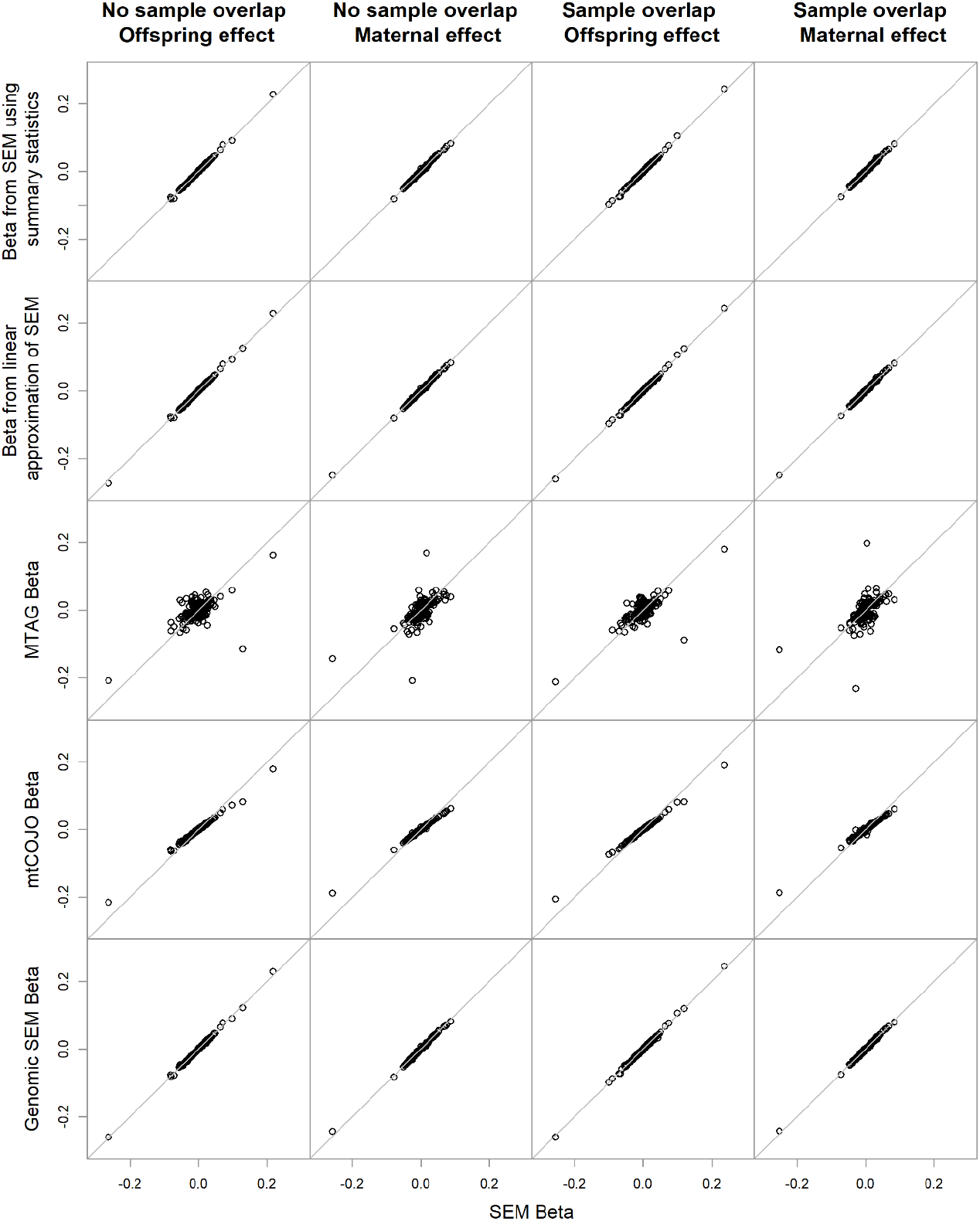
Comparison of the effect size estimates from the SEM using individual level data (x-axis) and the various different methods using the summary statistics from the GWAS of own and offspring birth weight (y-axis) for the 301 genome-wide significant SNPs from Warrington et al^3^. The columns summarize the results from the analysis including unique individuals in the GWAS of own and offspring birth weight for the offspring and maternal effect respectively, followed by the results from the analysis where there were overlapping samples in the GWAS of own and offspring birth weight for the offspring and maternal effect.

The comparison of the standard errors for the genetic effects for each of the 300 genetic variants are displayed in **Figure 3**. The standard errors for both the maternal and offspring effect are comparable to the SEM using individual level data when using Genomic SEM, both with and without sample overlap. They are also comparable using the linear approximation of the SEM when there is no sample overlap, but are slightly inflated relative to the SEM using individual level data when there is sample overlap that is not accounted for. This is expected as the standard error equations for the linear approximation would need to subtract twice the covariance between the maternal and offspring genetic effects when there is sample overlap. In contrast, the standard errors for the maternal and offspring effects are accurate using the SEM based on covariance matrices derived from the summary statistics when there is sample overlap that has been estimated using LD score regression and incorporated in the model, but they are under estimated when there is no sample overlap. This could be due to the slight sample overlap that was estimated by LD score regression (8,396 individuals were estimated to overlap both GWAS) and included in the SEM using covariance matrices but was not accounted for in the baseline SEM analysis using individual level data as there was no overlap. As expected due to the difference in purpose of the method, the standard errors estimated using MTAG and mtCOJO were smaller than those estimated by the SEM using individual level data for the maternal and offspring genetic effect, with and without sample overlap, with the largest difference for MTAG.

**Figure 3:**
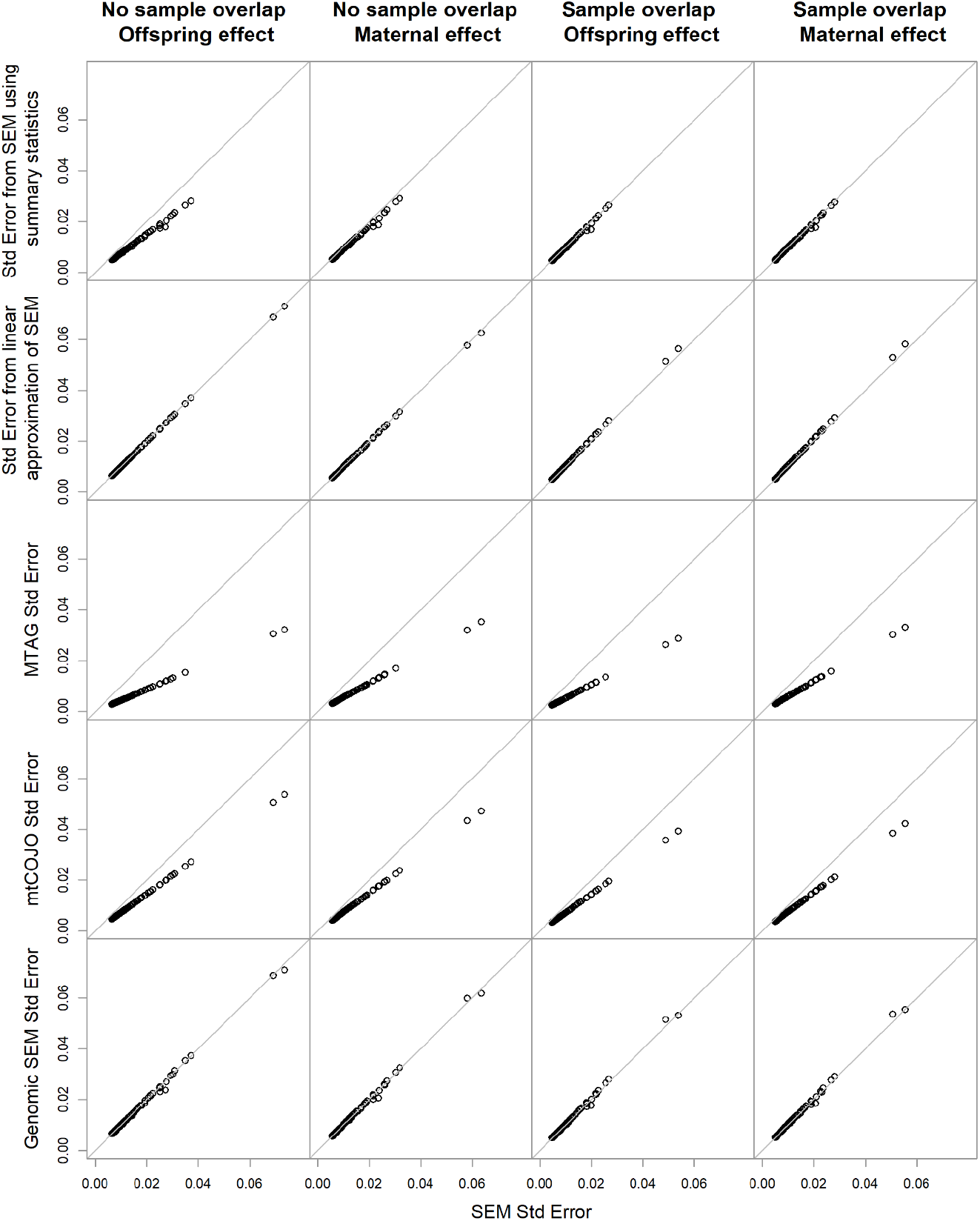
Comparison of the standard error from the SEM using individual level data (x-axis) and the various different methods using the summary statistics from the GWAS of own and offspring birth weight (y-axis) for the 301 genome-wide significant SNPs from Warrington et al^3^. The columns summarize the results from the analysis including unique individuals in the GWAS of own and offspring birth weight for the offspring and maternal genetic effect respectively, followed by the results from the analysis were there were overlapping samples in the GWAS of own and offspring birth weight for the offspring and maternal genetic effect.

We conducted heterogeneity tests for the 300 autosomal genome-wide significant SNPs between the SEM using individual level data and each of the summary statistics methods (**Supplementary Tables 1 and 2**). After Bonferoni correction for the number of SNPs with results, as we would expect, we identified 18 SNPs with significant heterogeneity between the MTAG estimates and the SEM for the offspring effect and 9 SNPs for the maternal effect (3 SNPs showed significant heterogeneity for both the maternal and offspring effect) when there was sample overlap between the GWAS of own and offspring birth weight (**Supplementary Table 1**). In contrast, were unable to detect significant heterogeneity for any SNPs using mtCOJO (offspring effect P_min_=0.039, maternal effect P_min_=0.083), the SEM based on covariance matrices derived from the summary statistics (offspring effect P_min_=0.700, maternal effect P_min_=0.585), the linear approximation of the SEM (offspring effect P_min_=0.707, maternal effect P_min_=0.595) or genomic SEM (offspring effect P_min_=0.762, maternal effect P_min_=0.758). Similar results were seen when there was no sample overlap between the GWAS of own and offspring birth weight (**Supplementary Table 2**).

### Comparison of computational time

The computational time was not influenced by sample overlap. MTAG, mtCOJO and the linear approximation of the SEM all took approximately between 30-60 minutes (see **Table 1** for precise computational time for each method). When running each chromosome in parallel, Genomic SEM took under four hours to complete. In contrast, the SEM based on covariance matrices derived from the summary statistics took over 60 hours to complete, indicating that it is much more computationally intensive than the other methods.

**Table 1:**
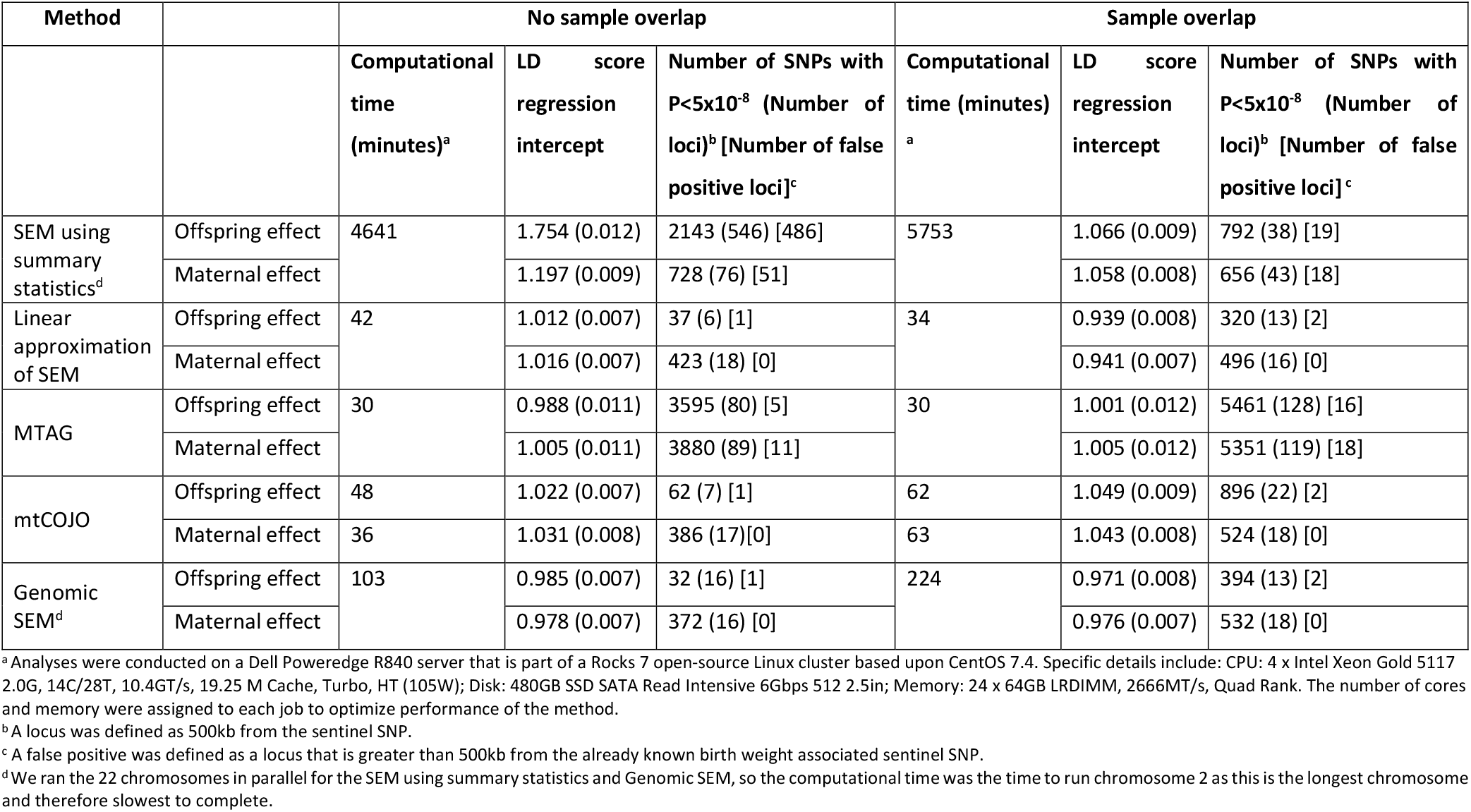
Summary of the methods used to derive maternal and offspring specific genetic effects using summary statistics from a GWAS of own birth weight and a GWAS of offspring birth weight. Computational time is used as a guide only to compare between methods; this will differ depending on the computing resources available for each analysis.

| Method |  | No sample overlap |  |  | Sample overlap |  |  |
| --- | --- | --- | --- | --- | --- | --- | --- |
| | | Computational time (minutes)<sup>a</sup> | LD score regression intercept | Number of SNPs with $P < 5 \times 10^{-8}$ (Number of loci)<sup>b</sup> [Number of false positive loci]<sup>c</sup> | Computational time (minutes)<sup>a</sup> | LD score regression intercept | Number of SNPs with $P < 5 \times 10^{-8}$ (Number of loci)<sup>b</sup> [Number of false positive loci]<sup>c</sup> |
| SEM using summary statistics<sup>d</sup> | Offspring effect | 4641 | 1.754 (0.012) | 2143 (546) [486] | 5753 | 1.066 (0.009) | 792 (38) [19] |
|  | Maternal effect |  | 1.197 (0.009) | 728 (76) [51] |  | 1.058 (0.008) | 656 (43) [18] |
| Linear approximation of SEM | Offspring effect | 42 | 1.012 (0.007) | 37 (6) [1] | 34 | 0.939 (0.008) | 320 (13) [2] |
|  | Maternal effect |  | 1.016 (0.007) | 423 (18) [0] |  | 0.941 (0.007) | 496 (16) [0] |
| MTAG | Offspring effect | 30 | 0.988 (0.011) | 3595 (80) [5] | 30 | 1.001 (0.012) | 5461 (128) [16] |
|  | Maternal effect |  | 1.005 (0.011) | 3880 (89) [11] |  | 1.005 (0.012) | 5351 (119) [18] |
| mtCOJO | Offspring effect | 48 | 1.022 (0.007) | 62 (7) [1] | 62 | 1.049 (0.009) | 896 (22) [2] |
|  | Maternal effect | 36 | 1.031 (0.008) | 386 (17) [0] | 63 | 1.043 (0.008) | 524 (18) [0] |
| Genomic SEM<sup>d</sup> | Offspring effect | 103 | 0.985 (0.007) | 32 (16) [1] | 224 | 0.971 (0.008) | 394 (13) [2] |
|  | Maternal effect |  | 0.978 (0.007) | 372 (16) [0] |  | 0.976 (0.007) | 532 (18) [0] |

<sup>a</sup> Analyses were conducted on a Dell Poweredge R840 server that is part of a Rocks 7 open-source Linux cluster based upon CentOS 7.4. Specific details include: CPU: 4 x Intel Xeon Gold 5117 2.0G, 14C/28T, 10.4GT/s, 19.25 M Cache, Turbo, HT (105W); Disk: 480GB SSD SATA Read Intensive 6Gbps 512 2.5in; Memory: 24 x 64GB LRDIMM, 2666MT/s, Quad Rank. The number of cores and memory were assigned to each job to optimize performance of the method.
<sup>b</sup> A locus was defined as 500kb from the sentinel SNP.
<sup>c</sup> A false positive was defined as a locus that is greater than 500kb from the already known birth weight associated sentinel SNP.
<sup>d</sup> We ran the 22 chromosomes in parallel for the SEM using summary statistics and Genomic SEM, so the computational time was the time to run chromosome 2 as this is the longest chromosome and therefore slowest to complete.

### Evidence of inflation of test statistics across the genome

Manhattan plots and Q-Q plots for each of the conditional GWAS are presented in **Supplementary Figures 4-23**. The LD score regression intercepts and number of genome-wide significant (P<5×10^-8^) SNPs are presented in **Table 1**. Inflation in the test statistics was detected for the SEM based on covariance matrices derived from the summary statistics when there was no sample overlap (LD score regression intercepts: offspring = 1.754, maternal = 1.197). This resulted in a larger number of variants/loci being identified as genome-wide significant (**Table 1**). This appears to predominantly be driven by the underestimation of the standard error (as seen in **Figure 3**), which is most prominent for SNPs with lower minor allele frequency or SNPs where the maternal and offspring genetic effect are going in opposite directions. Deflation in the test statistics was detected for the linear approximation of the SEM when there was sample overlap (LD score regression intercepts: offspring = 0.939, maternal = 0.941). This is expected as the standard errors are slightly larger because the equations do not account for sample overlap. The LD score regression intercepts for the other methods ranged between 0.971 and 1.066, indicating that there was not much genome-wide inflation in the test statistics.

We defined a birth weight associated locus as being 500kb from a previously identified birth weight associated sentinel SNP^24^. Although a large number of birth weight associated loci were detected using some of the methods (5-546 loci; **Table 1**), the majority of them have been previously associated with birth weight (which we assume are true positives). The SEM using summary statistics detected a large number of what are presumably false positives (those loci that have not previously been associated with birth weight in far larger samples of individuals^24^), particularly when there is no sample overlap, which is in line with the inflated LD score intercepts. MTAG also detected a substantial number of false positives, whereas the three other methods only detected up to three false positives.

### Results from simulations

The simulated data showed substantial deflation in the test statistics under the null for both the maternal and offspring specific genetic effects on birth weight, with or without sample overlap, when using mtCOJO (**Supplementary Figures 24**). This was driven by both an underestimation of the effect size using mtCOJO and an overestimation of the standard error in comparison to the approximation of the SEM which is conditioning on the genotype rather than the phenotype (**Supplementary Figures 25 and 26**). This is in contrast to the underestimation of the standard error when there is a true effect, as seen in **Figure 3** for the previously known birth weight associated loci. Therefore, it appears that conditioning on the phenotype using mtCOJO is a good approximation of conditioning on the genotype when the locus is associated with the outcome, however it is a poor approximation under the null.

### Fertility GWAS

Through the methods comparison, we have shown that Genomic SEM out performs the other methods in terms its ability to accurately estimate conditional effect sizes and standard errors at individual genetic variants and its ability to account for sample overlap appropriately. It is also a highly flexible method, allowing the estimation of other genetic effects, such as paternal specific effects. To illustrate application of this method and how it can be extended to simultaneously estimate conditional maternal, paternal and offspring genetic effects, we applied it to fertility data from the UK Biobank.

Results from the unconditional GWAS analysis conducted in BOLT-LMM of the number of children fathered, the number of children mothered and the number of siblings can be visualised in **Supplementary Figure 27**. We conducted two separate analyses in Genomic SEM; firstly we calculated only the female and sibling specific effects using the GWAS results from number of children mothered and number of siblings (**Supplementary Figure 28**), and secondly we utilized all three GWAS to calculate the female, male and sibling specific effects (**Figure 4**). Using LD score regression^21^, we estimated the genetic correlations between the unconditional and conditional GWAS. There was a strong genetic correlation between the number of children mothered (unconditional) and the female-specific effect on fertility (analysis one: r_g_=0.941, SE=0.006; analysis two: r_g_=0.932, SE=0.008) and similarly between the number of children fathered and the male-specific effect on fertility (analysis two: r_g_=0.904, SE=0.013). In contrast, the genetic correlation was weaker between the number of siblings and the sibling specific effect, particularly once male specific effects were incorporated (analysis one: r_g_=0.712, SE=0.025; analysis two: r_g_=0.173, SE=0.058). We also used LD score regression to estimate the genetic correlation between the conditional GWAS for male and female fertility (r_g_=0.871, SE=0.023), male fertility and sibling effects (r_g_= −0.826, SE=0.023) and female fertility and sibling effects (r_g_= −0.812, SE=0.019).

**Figure 4:**
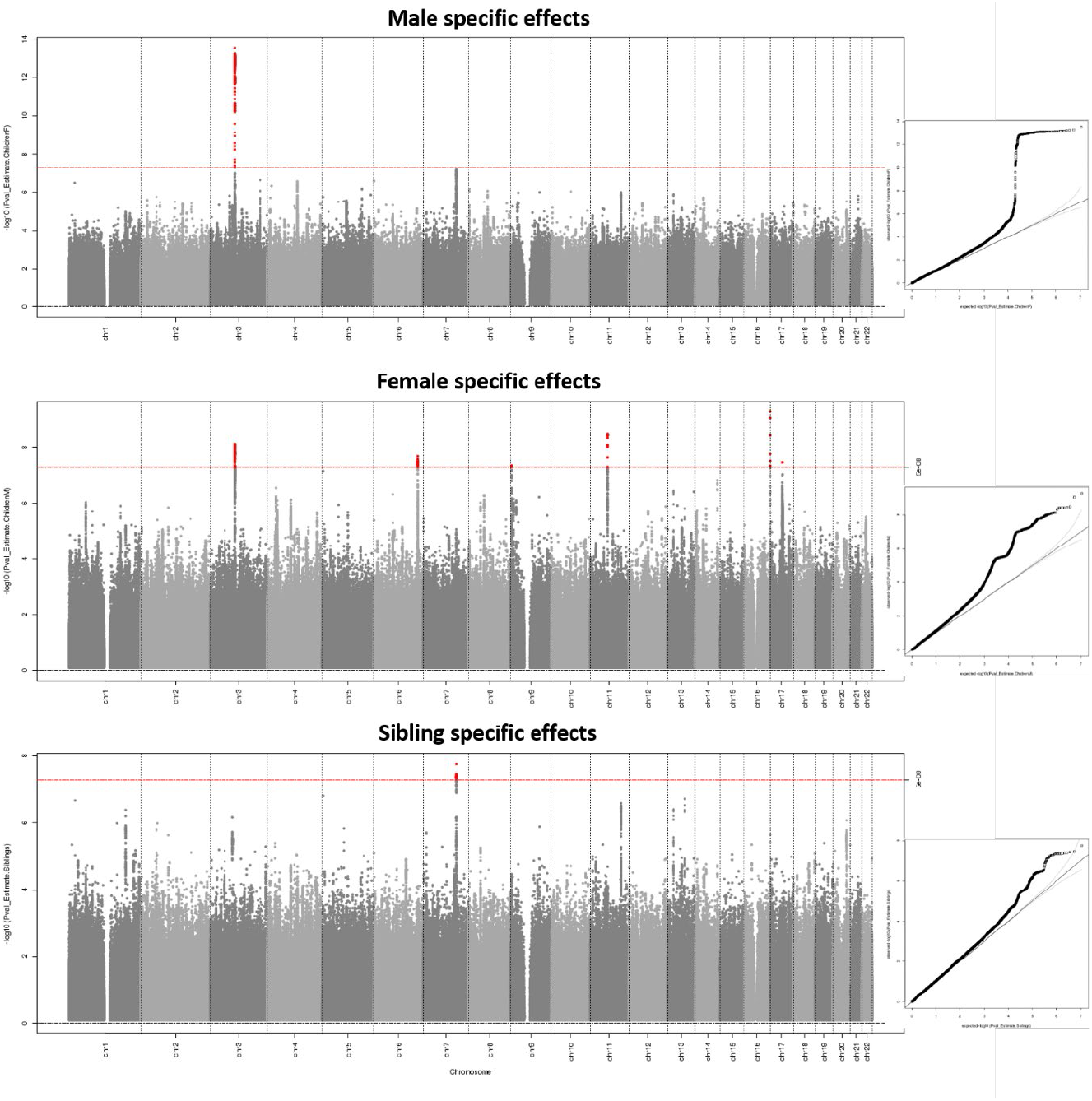
Manhattan plot and quantile-quantile (Q-Q) plot for the fertility GWAS estimating male, female and sibling specific genetic effects using Genomic SEM. 237,768 women from the UK Biobank contributed to the unconditional GWAS of the number of children mothered, 199,570 men contributed to the GWAS of the number of children fathered and 430,466 individuals contributed to the GWAS of the number of siblings (see Supplementary Figure 27 for the Manhattan plots of the unconditional GWAS). The two-sided association P-value, on the −log10 scale, obtained from Genomic SEM for each of the SNPs (y-axis) was plotted against the genomic position (NCBI Build 37; x-axis). Association signals that reached genome-wide significance (P < 5×10^-8^) are shown in red. In the Q-Q plots, the black dots represent observed P-values and the grey line represents expected P-values under the null distribution. The SNP heritability, estimated using LD score regression, was 0.033 (SE=0.003) for male fertility, 0.042 (SE=0.003) for female fertility and 0.012 (SE=0.001) for sibling specific effects.

Both male and female fertility from the conditional analysis were negatively genetically correlated with years of education (male r_g_=-0.17, SE=0.03, P=7×10^-8^; female r_g_=-0.20, SE=0.03, P=3×10^-13^) and positively genetically correlated with risk taking behaviours (male r_g_=0.27, SE=0.05, P=1×10^-7^; female r_g_=0.17, SE=0.04, P=6×10^-5^; **Supplementary Figure 29**), whereas sibling effects were not correlated with years of education or risk taking behaviours. Additionally, male fertility was negatively genetically correlated with autism spectrum disorder (r_g_=-0.24, SE=0.07, P=6×10^-4^). Subjective wellbeing was also genetically correlated with fertility; positively correlated with male fertility (r_g_=0.23, SE=0.06, P=7×10^-5^) and negatively correlated with sibling effects (r_g_=-0.23, SE=0.06, P=3×10^-4^).

The results from analysis one estimating the female and sibling specific effects only on fertility, where we identified four loci (P<5×10^-8^) associated with the number of children mothered, after conditioning on the number of siblings, and one locus associated with the number of siblings, conditional on the number of children mothered. When we extended the Genomic SEM model to estimate female, male and sibling specific genetic effects in analysis two, we identified six loci associated with maternal-specific effects, one locus associated with paternal-specific effects (in the same region as one of the maternal-specific loci) and one locus associated with sibling-specific effects (**Figure 4**). After conditioning on male fertility, the locus associated with a sibling specific effect in the female/sibling only analysis attenuated slightly (P=3.6×10^-5^), even though it is a different locus to the one identified on chromosome 3 for the male and female specific effects. The full results for each of these genome-wide significant loci are presented in **Supplementary Table 3**. Interestingly, a number of the genes nearest to our genome-wide significant loci have previously been associated with age at first sexual intercourse (*ESR1, CADM2*), number of sexual partners (*CADM2*), educational attainment (*ESR1, TUBB3, MC1R, CADM2, MDFIC*) and risk-taking behaviour (*MDFIC*).

## Discussion

We compared five different statistical methods to estimate maternal and offspring specific genetic effects on an offspring outcome using summary statistics from GWAS and have shown that Genomic SEM outperforms the other methods in terms of estimation of the effect size and standard error, ability to account for sample overlap appropriately, and flexibility to estimate other genetic effects such as a paternal specific effect. It was more time consuming than several of the other methods; however, running the chromosomes in parallel allowed the GWAS to be completed in under four hours. Additionally, we detected some deflation in the test statistics that could have been due to the use of the stricter version of genomic control that was implemented in the version of the software used for this analysis; this has been relaxed in more recent releases. Subsequently, we used Genomic SEM to identify the genetic loci associated with male and female fertility, after adjusting for sibling genetic effects, and identified seven loci, one of which was novel. We note that the focus of this paper is on partitioning the genetic effect into indirect parental and direct offspring genetic components, and not necessarily to identify novel loci associated with the traits of interest.

There were several limitations with using summary statistics in the SEM, which resulted in inflation in the test statistics. Firstly, we estimated the sample overlap using LD score regression^21^, which was overestimated in both of our analyses with and without sample overlap. This overestimation has been described previously when there is population stratification^25^, and the authors suggest a modified formula to calculate the overlap. We did not use this modified formula in the current analysis as we and others have previously used LD Score regression to estimate sample overlap and we wanted to get an idea of how this would perform^24,26^. It could also be due to cryptic relatedness between the GWAS; for example, there might be close relatives in both GWAS of own and offspring birth weight that are adding to this overestimation. Secondly, estimates of the maternal and offspring specific genetic effects can vary dramatically if the phenotypic correlation between the maternal and offspring phenotype is mis-specified. Given we had access to the phenotypic data that was used for the GWAS analysis, we were able to obtain a good estimate of the phenotypic correlation; however, if using publically available GWAS results this would be more difficult to estimate accurately. Thirdly, this method assumes that the effect size estimates from the unadjusted GWAS are estimated accurately and does not account for the standard errors. Therefore, for low frequency genetic variants that have large standard errors, we saw the method performed poorly. We also found the method performed poorly for a subset of genetic variants where the maternal specific genetic effect on the offspring outcome went in the opposite direction to the offspring specific genetic effect, particularly when there was no sample overlap between the unadjusted GWAS. However, as we showed in the initial paper describing the SEM^9^, including some raw data in addition to the covariance matrices estimated from the summary statistics improves estimation of the maternal and offspring specific genetic effects.

The linear approximation of the SEM assumes no sample overlap, so performed well when there was no overlap; however, the standard errors were overestimated when there was sample overlap, deflating the test statistics. The formula for estimating the standard error of the maternal and offspring specific effects could be extended to account for any sample overlap, but it would need to rely on LD score regression to estimate the sample overlap which has issues as described previously. Otherwise, this method performed well in terms of accurately estimating the maternal and offspring specific effects and was one of the fastest methods to perform the conditional analysis.

Given that MTAG is not designed to estimate maternal and offspring specific effects at individual loci, it was not surprising that it performed the poorest at partitioning the genetic effect into maternal and offspring components out of all the methods in our comparison. This is because it is estimating an overall genetic effect on the offspring’s outcome, and not partitioning it into maternal and offspring components. In other words, the effect estimate for the offspring specific effect at an individual genetic variant was equivalent to the maternal specific effect at that same variant. This is the most powerful approach of those compared if you want to detect novel loci for an offspring outcome, but one of the other methods would subsequently be required to partition the genetic effect into maternal and offspring components.

Finally, mtCOJO was developed to condition the outcome on one or more covariate phenotypes; whereas the other methods we compare are equivalent to conditioning the outcome on the genotype. For example, to estimate the offspring specific genetic effect mtCOJO is conditioning the genetic effect on offspring birth weight rather than maternal genotype. When the genetic variant is associated with the outcome, conditioning on the phenotype in mtCOJO seems to proxy conditioning on the genotype well and reduces the standard error (increasing the statistical power to detect an association). However, as seen in our simulations, when the genetic variant is not associated with the outcome, conditioning on the phenotype does not proxy conditioning on the genotype well and the standard error is overestimated.

We performed the first GWAS partitioning the genetic effect into male and female fertility specific genetic effects and a sibling specific effect. The heritability of number of children ever born has been estimated to be between 0.24-0.43^27^, and the variance explained by common genetic variants (SNP based heritability) is estimated to be approximately 10%^28^. Although there have been two GWAS previously conducted on number of children born^23,29^, no study to date has estimated the conditional male, female and sibling genetic effects at individual genetic loci. Both previous studies observed significant genetic correlations for the number of children between men and women (Barban and colleagues^23^: r_g_=0.97, SE=0.095; Mathieson and colleagues^29^: r_g_=0.74, 95%CI=0.66-0.82), which our findings from the conditional analysis are consistent with (r_g_=0.871, SE=0.023). We also identified a strong negative genetic correlation between both male/female fertility and sibling effects; this is likely to be due to a technical artefact of the analyses as described by Wu and colleagues^30^. We replicate the negative genetic correlation between years of education and fertility described in Barban *et al*^23^. Furthermore, we found a positive genetic correlation between risk taking behaviour and both male and female fertility, showing that having more increasing alleles for the number of children is associated with a higher genetic risk for partaking in risk taking behaviours. Due to our ability to partition the genetic effect, we were also able to identify a negative genetic correlation between male fertility and autism, indicating that fathers at genetically increased risk of autism are more likely to have fewer children. This relationship between fertility and autism, particularly in males, has previously been shown in a large population based study in Sweden using patients with various psychiatric disorders and their unaffected siblings^31^. Interestingly, we identified a positive genetic correlation between male fertility and subjective well-being and a negative genetic correlation with the sibling effect. We identified six of the previous 28 statistically independent loci for fertility, and one of the 16 loci for childlessness^29^. In addition, we identified a novel locus on chromosome 9 that is associated with female fertility. This locus is near *RFX3*, which harbours genetic variants that have previously been associated with smoking initiation. Our genetic correlation analysis shows a positive genetic correlation between both male and female fertility and smoking initiation, however it does not meet our multiple testing threshold.

In conclusion, when estimating maternal and offspring (and paternal) specific genetic effects on an offspring outcome using GWAS summary statistics, we recommend using Genomic SEM.

## Acknowledgements

This research was carried out at the Translational Research Institute, Woolloongabba, QLD 4102, Australia. The Translational Research Institute is supported by a grant from the Australian Government. This study has been conducted using the UK Biobank Resource under Application Number 53641. D.M.E. is supported by an Australian National Health and Medical Research Council Senior Research Fellowship (1137714) and this work was supported by a Australian National Health and Medical Research Council Project Grant (1157714) and Ideas Grant (1183074).

