## Supplementary Material for "Estimating direct and indirect genetic effects on offspring phenotypes using genome-wide summary results data: a comparison of multivariate methods"

**Supplementary Figure 1: Diagram of the structural equation model (SEM) for estimating maternal and offspring genetic effects on birth weight.** The three observed variables (in squares) are the birth weight of the individual (BW), the birth weight of their offspring ($BW_O$) and the genotype of the individual (SNP). The latent variables (in circles) are the genotypes for the individual's mother ($G_G$) and the genotype of the individual's first offspring ($G_O$). The total variance of the latent genotypes for the individual's mother ($G_G$) and offspring ($G_O$) and for the observed SNP variable is set to Φ (i.e., variance($G_G$) = Φ, variance (SNP) = 0.75Φ + 0.25Φ, variance ($G_O$) = 0.75Φ + 0.25Φ). $\hat{\beta}_{m_{adj}}$ and $\hat{\beta}_{o_{adj}}$ path coefficients refer to maternal and offspring effects respectively. The residual error terms for the birth weight of the individual and their offspring are represented by ε and $\varepsilon_O$ respectively and we estimate the variance of both of these terms in the SEM. The covariance between residual genetic and environmental sources of variation is given by $\rho$. A) is used to model the subset of individuals with complete data. B) is used to model the subset of genotyped individuals who report their own birth weight, but not their offspring's birth weight. Genotyped males who report their own birth weight (but not their offspring's) can be incorporated into this part of the model. C) is used to model the subset of genotyped individuals who report their offspring's birth weight, but not their own. These three models are fit to the three subsets of data that contain the various patterns of missingness, and then the likelihoods from each model are combined. In the analysis with no sample overlap, the likelihoods from model B and C were combined and therefore no correlation between the own and offspring birthweight was estimated.

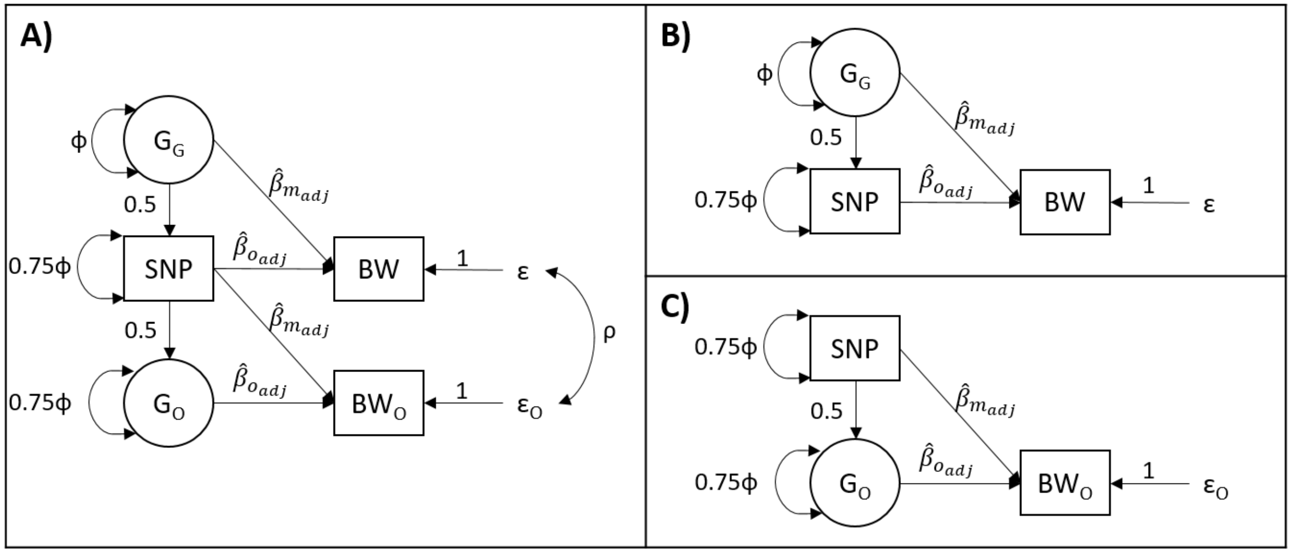

**Supplementary Figure 2: Diagram of the genomic structural equation model (Genomic SEM) for estimating conditional genetic effects on birthweight.** The two 'observed variables' (in squares) are the summary results statistics from the genome-wide association study (GWAS) of birth weight of the individual and from the GWAS of the birth weight of their offspring. The latent variables (in circles) are the constructs that we are estimating the genetic effects of. $\hat{\beta}_{m_{adj}}$ and $\hat{\beta}_{o_{adj}}$ path coefficients refer to maternal and offspring effects respectively. The residual error terms for the birth weight of the individual and their offspring are represented by $\phi_o$ and $\phi_m$ respectively. Finally, ρ represents the genetic covariance between the latent factors, which can be estimated using Genomic SEM (it was not estimated in our model, hence the dotted double arrow).

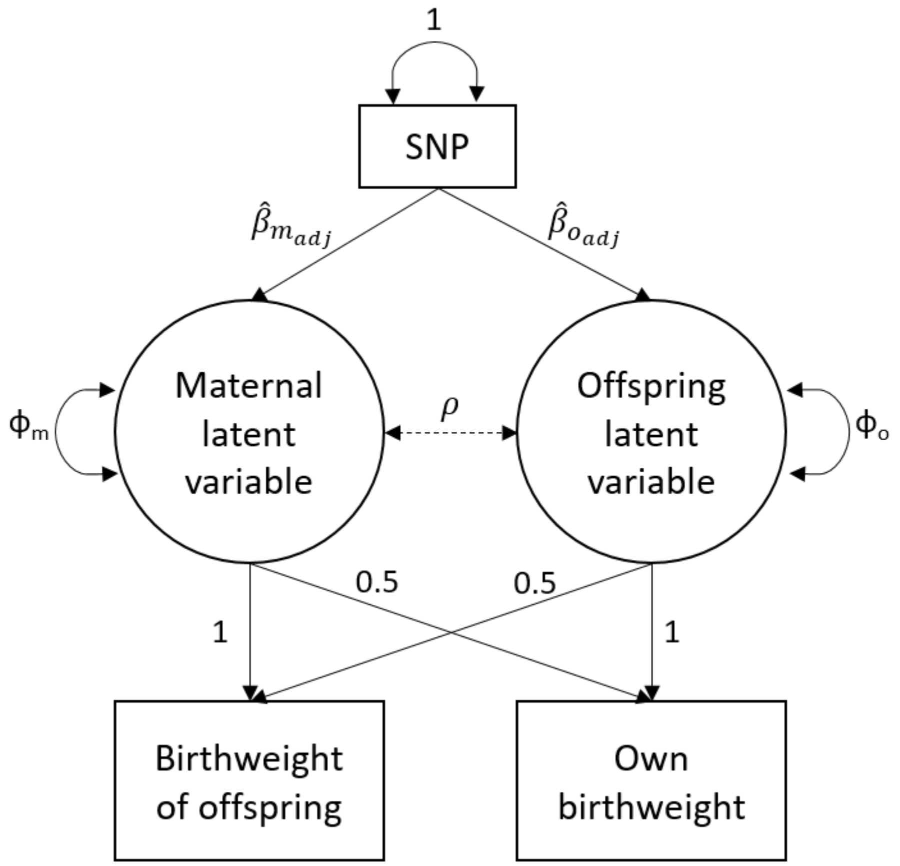

**Supplementary Figure 3: Diagram of the genomic structural equation model (Genomic SEM) for estimating conditional genetic effects on fertility.** A) is the model that is similar to the birth weight analysis with just the number of siblings and the number of children mothered. B) is the full model incorporating data on the number of children fathered.

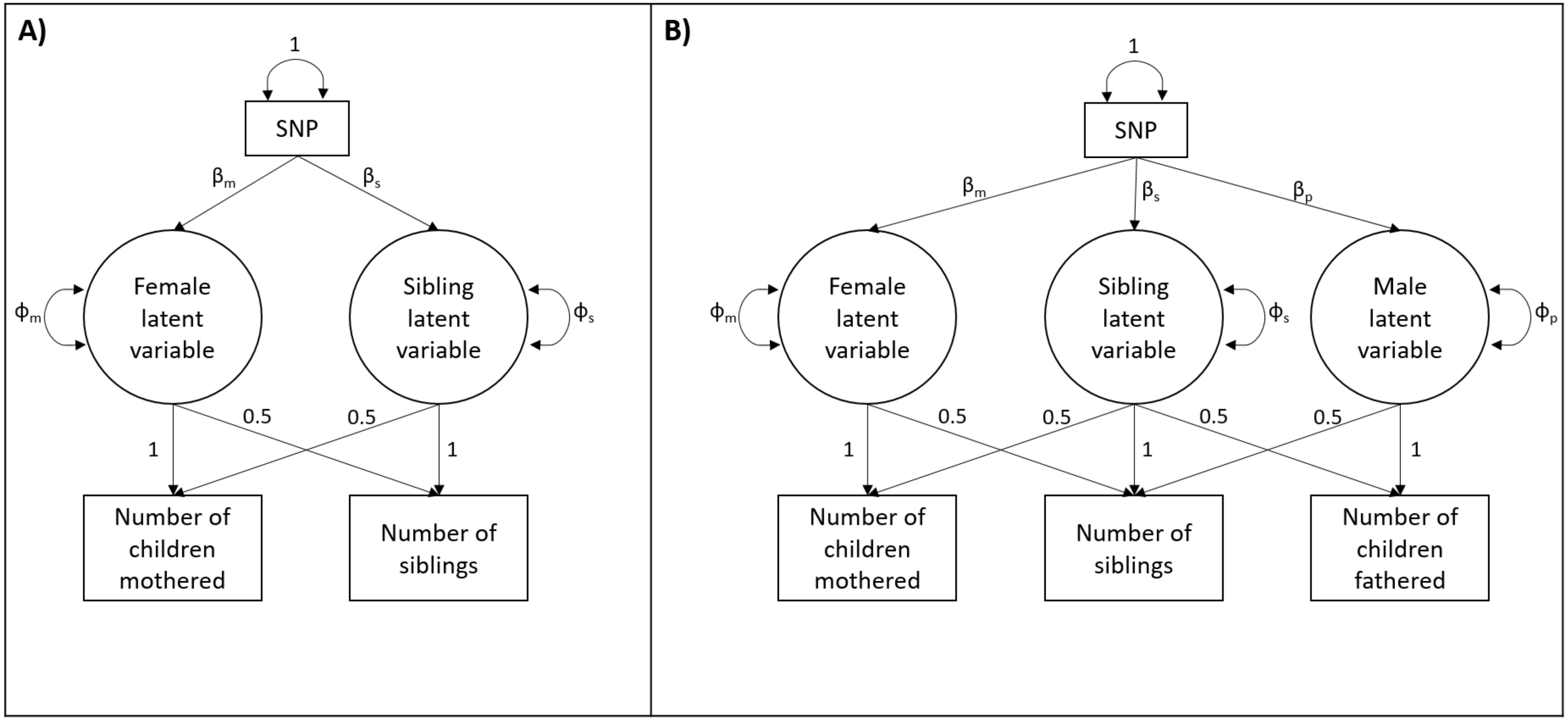

**Supplementary Figure 4: Manhattan plot and quantile-quantile (Q-Q) plot for the offspring specific effect estimated using the structural equation model (SEM) with summary statistics and no sample overlap between the GWAS of own and offspring birth weight.** The two-sided association P-value, on the –log10 scale, obtained from the SEM for each of the SNPs (y-axis) was plotted against the genomic position (NCBI Build 37; x-axis). Association signals that reached genome-wide significance ($P < 5x10^{-8}$) are shown in red if they are novel and turquoise if they have been previously reported[1]. In the Q-Q plots, the black dots represent observed P-values and the grey line represents expected P-values under the null distribution. The green dots represent observed P-values after excluding the previously identified signals[1].

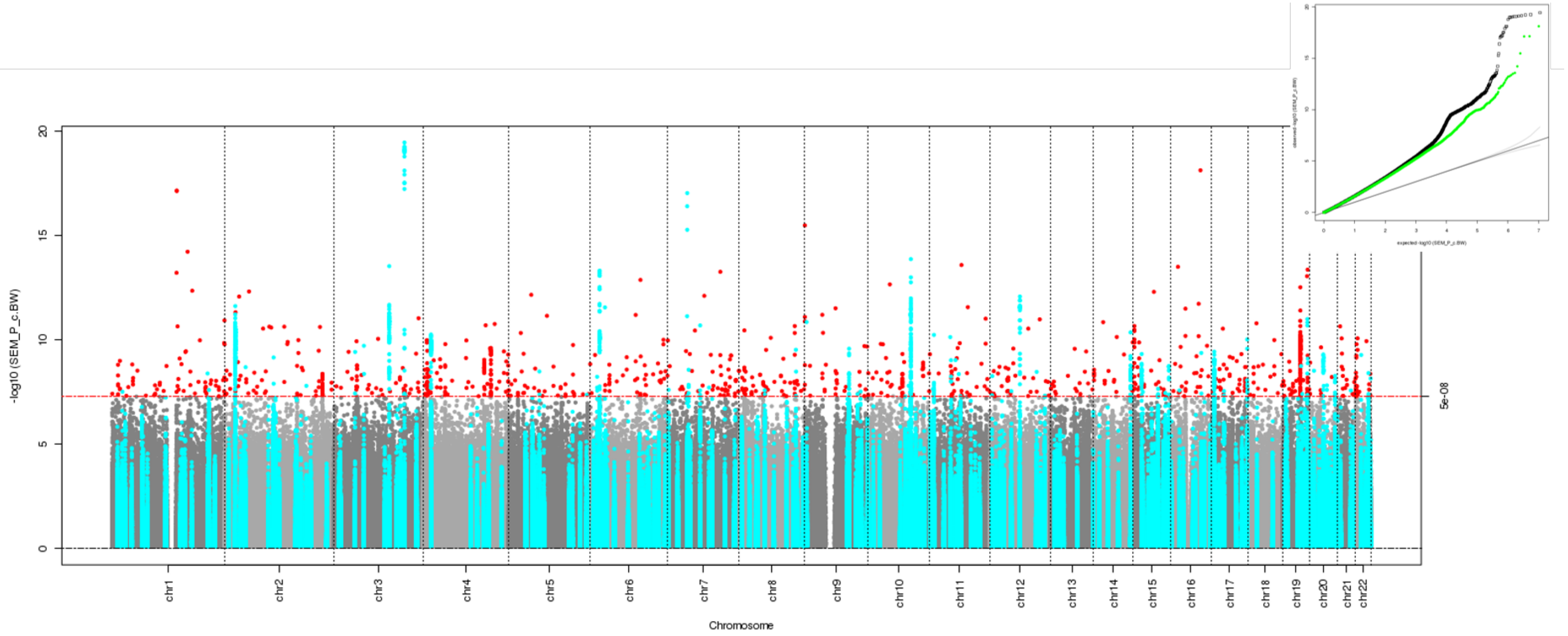

**Supplementary Figure 5: Manhattan plot and quantile-quantile (Q-Q) plot for the offspring specific effect estimated using the linear approximation of the structural equation model (SEM) and no sample overlap between the GWAS of own and offspring birth weight.** The two-sided association P-value, on the –log10 scale, obtained from the linear approximation of the SEM for each of the SNPs (y-axis) was plotted against the genomic position (NCBI Build 37; x-axis). Association signals that reached genome-wide significance ($P < 5x10^{-8}$) are shown in red if they are novel and turquoise if they have been previously reported[1]. In the Q-Q plots, the black dots represent observed P-values and the grey line represents expected P-values under the null distribution. The green dots represent observed P-values after excluding the previously identified signals[1].

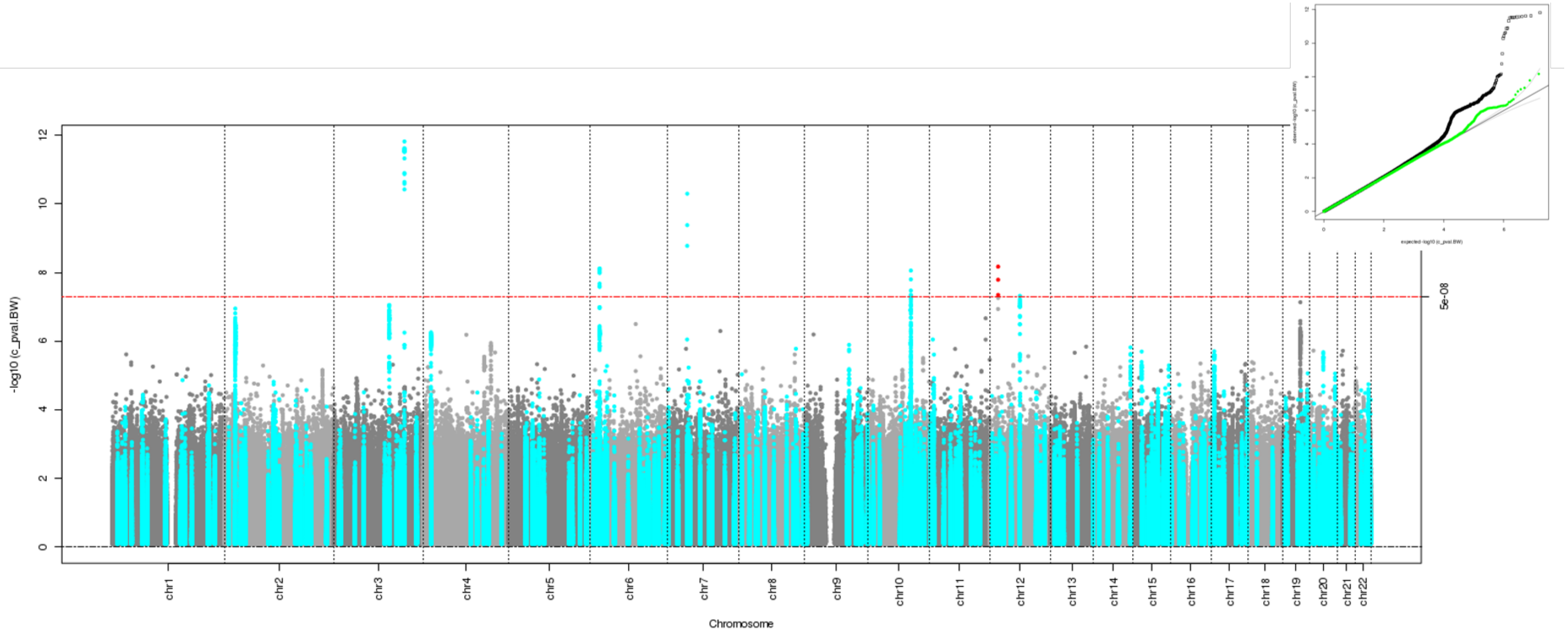

**Supplementary Figure 6: Manhattan plot and quantile-quantile (Q-Q) plot for the offspring specific effect estimated using MTAG[2] and no sample overlap between the GWAS of own and offspring birth weight.** The two-sided association P-value, on the –log10 scale, obtained from MTAG for each of the SNPs (y-axis) was plotted against the genomic position (NCBI Build 37; x-axis). Association signals that reached genome-wide significance ($P < 5x10^{-8}$) are shown in red if they are novel and turquoise if they have been previously reported[1]. In the Q-Q plots, the black dots represent observed P-values and the grey line represents expected P-values under the null distribution. The green dots represent observed P-values after excluding the previously identified signals[1].

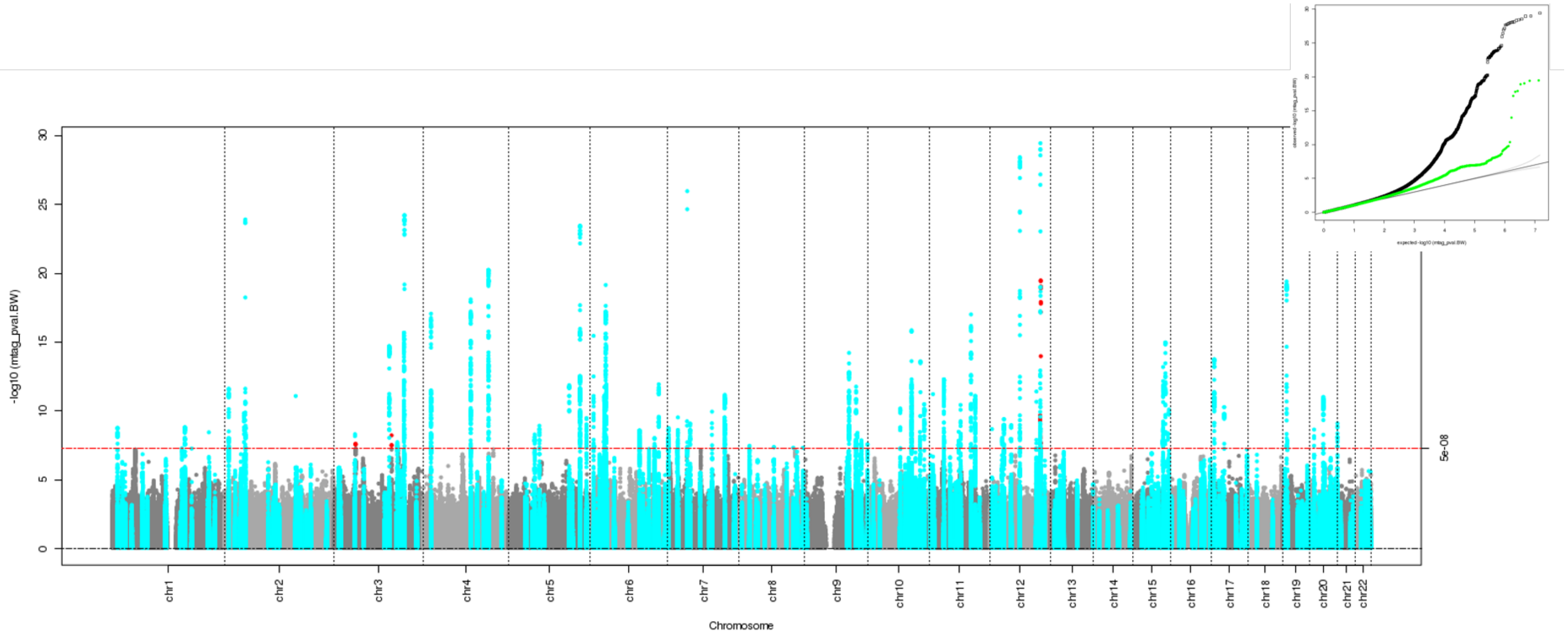

**Supplementary Figure 7: Manhattan plot and quantile-quantile (Q-Q) plot for the offspring specific effect estimated using mtCOJO[3] and no sample overlap between the GWAS of own and offspring birth weight.** The two-sided association P-value, on the –log10 scale, obtained from mtCOJO for each of the SNPs (y-axis) was plotted against the genomic position (NCBI Build 37; x-axis). Association signals that reached genome-wide significance ($P < 5x10^{-8}$) are shown in red if they are novel and turquoise if they have been previously reported[1]. In the Q-Q plots, the black dots represent observed P-values and the grey line represents expected P-values under the null distribution. The green dots represent observed P-values after excluding the previously identified signals[1].

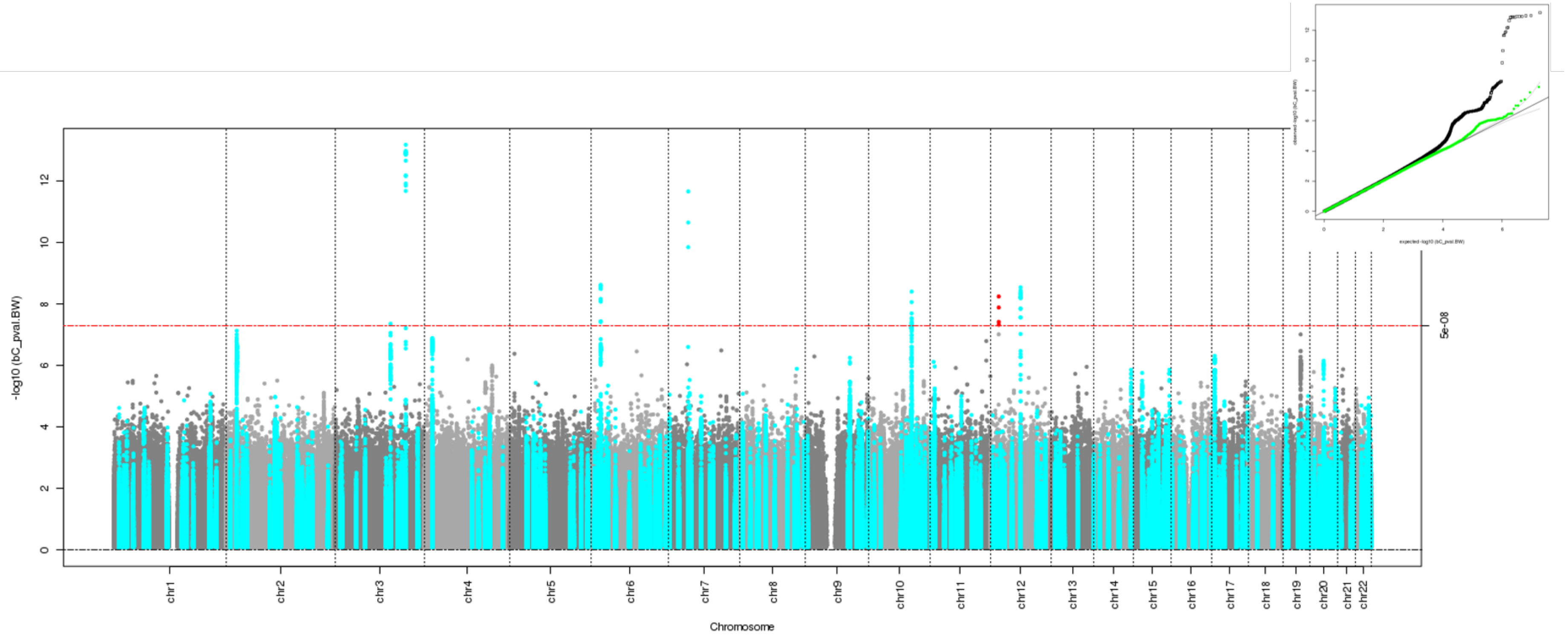

**Supplementary Figure 8: Manhattan plot and quantile-quantile (Q-Q) plot for the offspring specific effect estimated using Genomic SEM[4] and no sample overlap between the GWAS of own and offspring birth weight.** The two-sided association P-value, on the –log10 scale, obtained from Genomic SEM for each of the SNPs (y-axis) was plotted against the genomic position (NCBI Build 37; x-axis). Association signals that reached genome-wide significance ($P < 5x10^{-8}$) are shown in red if they are novel and turquoise if they have been previously reported[1]. In the Q-Q plots, the black dots represent observed P-values and the grey line represents expected P-values under the null distribution. The green dots represent observed P-values after excluding the previously identified signals[1].

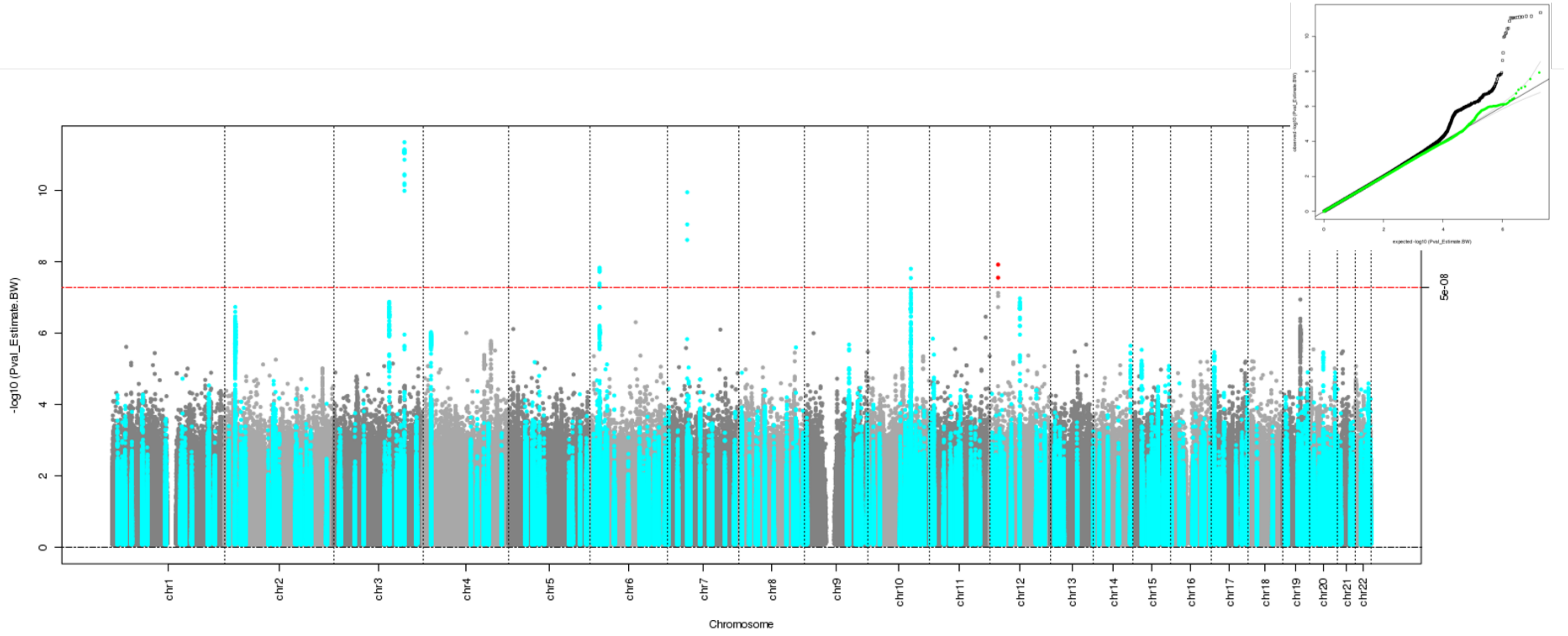

**Supplementary Figure 9: Manhattan plot and quantile-quantile (Q-Q) plot for the offspring specific effect estimated using the structural equation model (SEM) with summary statistics and sample overlap between the GWAS of own and offspring birth weight.** The two-sided association P-value, on the –log10 scale, obtained from the SEM for each of the SNPs (y-axis) was plotted against the genomic position (NCBI Build 37; x-axis). Association signals that reached genome-wide significance ($P < 5x10^{-8}$) are shown in red if they are novel and turquoise if they have been previously reported[1]. In the Q-Q plots, the black dots represent observed P-values and the grey line represents expected P-values under the null distribution. The green dots represent observed P-values after excluding the previously identified signals[1].

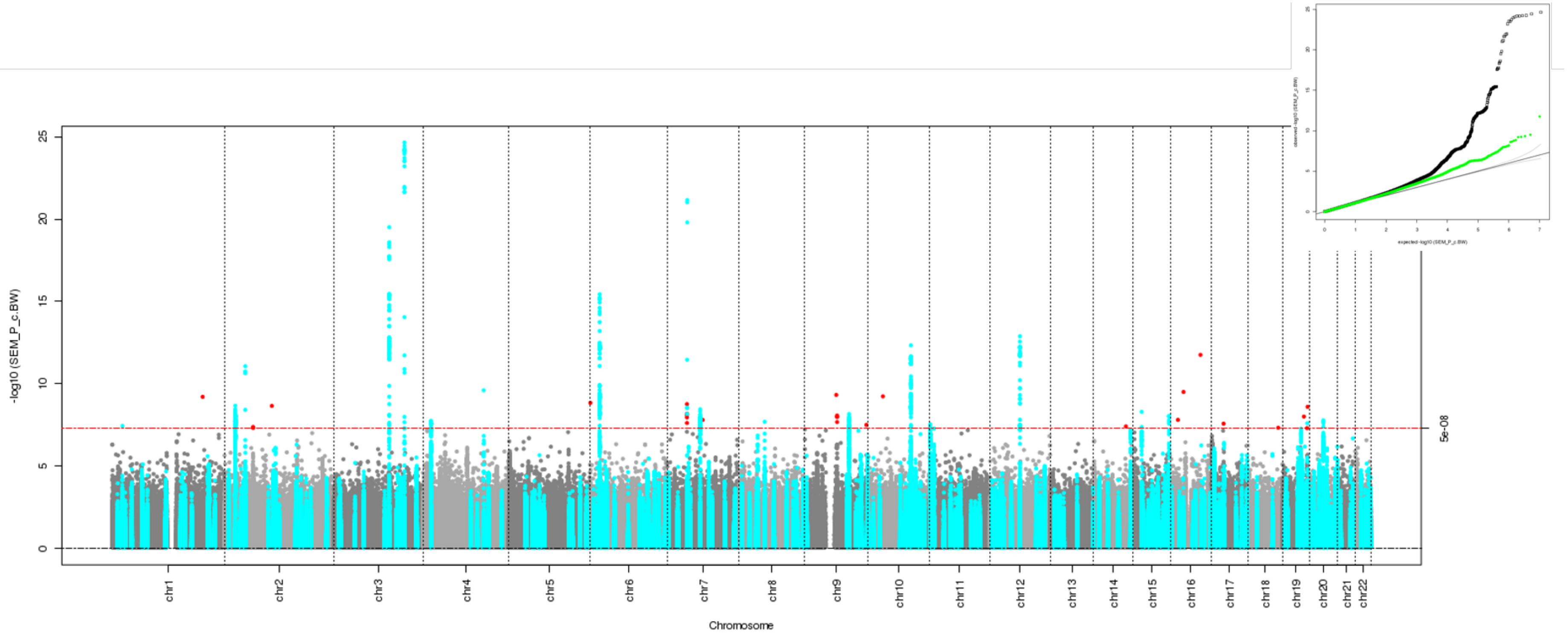

**Supplementary Figure 10: Manhattan plot and quantile-quantile (Q-Q) plot for the offspring specific effect estimated using the linear approximation of the structural equation model (SEM) and sample overlap between the GWAS of own and offspring birth weight.** The two-sided association P-value, on the –log10 scale, obtained from the linear approximation of the SEM for each of the SNPs (y-axis) was plotted against the genomic position (NCBI Build 37; x-axis). Association signals that reached genome-wide significance ($P < 5x10^{-8}$) are shown in red if they are novel and turquoise if they have been previously reported[1]. In the Q-Q plots, the black dots represent observed P-values and the grey line represents expected P-values under the null distribution. The green dots represent observed P-values after excluding the previously identified signals[1].

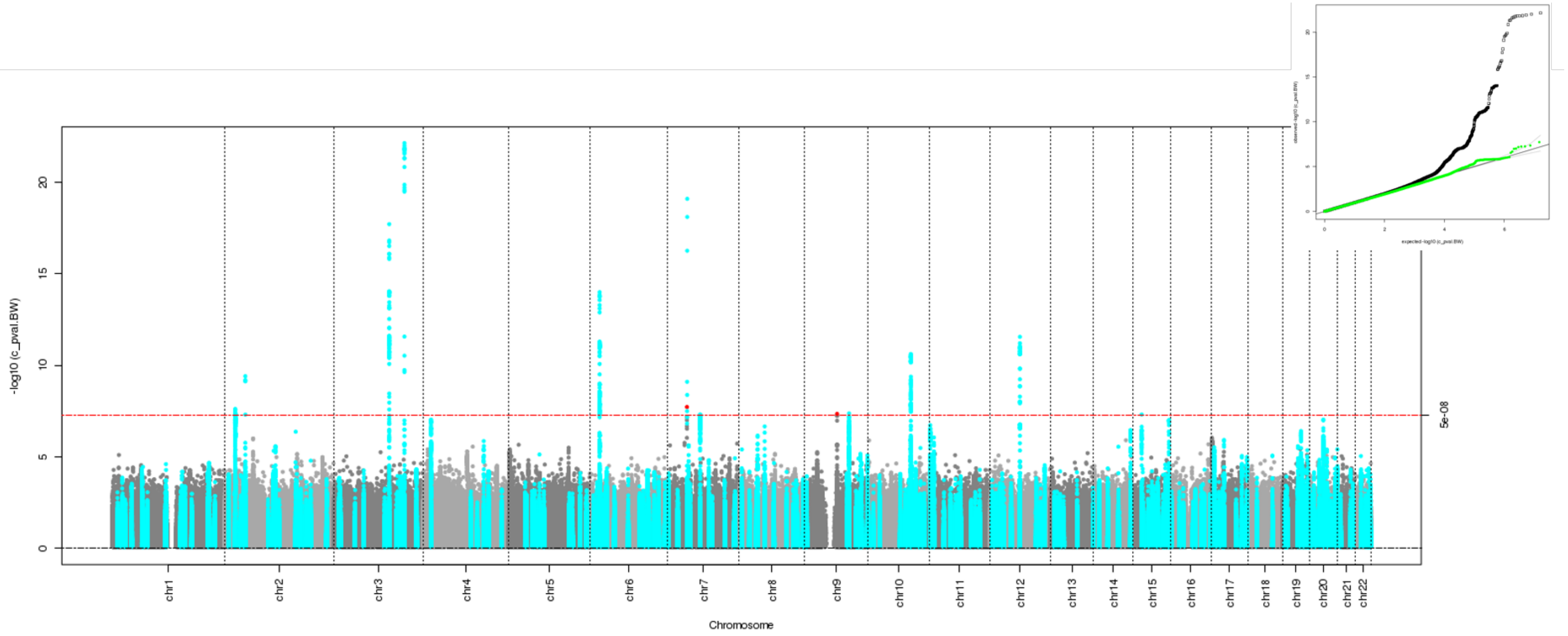

**Supplementary Figure 11: Manhattan plot and quantile-quantile (Q-Q) plot for the offspring specific effect estimated using MTAG[2] and sample overlap between the GWAS of own and offspring birth weight.** The two-sided association P-value, on the –log10 scale, obtained from MTAG for each of the SNPs (y-axis) was plotted against the genomic position (NCBI Build 37; x-axis). Association signals that reached genome-wide significance ($P < 5x10^{-8}$) are shown in red if they are novel and turquoise if they have been previously reported[1]. In the Q-Q plots, the black dots represent observed P-values and the grey line represents expected P-values under the null distribution. The green dots represent observed P-values after excluding the previously identified signals[1].

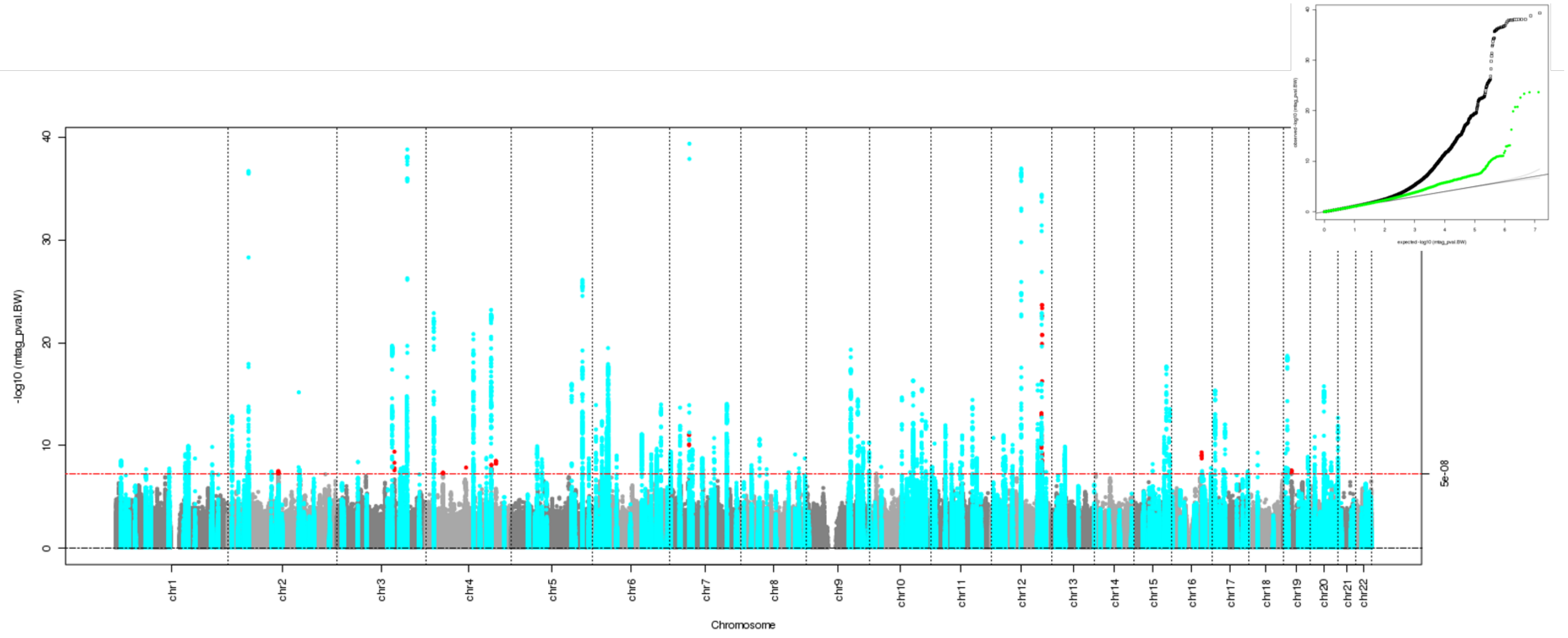

**Supplementary Figure 12: Manhattan plot and quantile-quantile (Q-Q) plot for the offspring specific effect estimated using mtCOJO[3] and sample overlap between the GWAS of own and offspring birth weight.** The two-sided association P-value, on the –log10 scale, obtained from mtCOJO for each of the SNPs (y-axis) was plotted against the genomic position (NCBI Build 37; x-axis). Association signals that reached genome-wide significance ($P < 5x10^{-8}$) are shown in red if they are novel and turquoise if they have been previously reported[1]. In the Q-Q plots, the black dots represent observed P-values and the grey line represents expected P-values under the null distribution. The green dots represent observed P-values after excluding the previously identified signals[1].

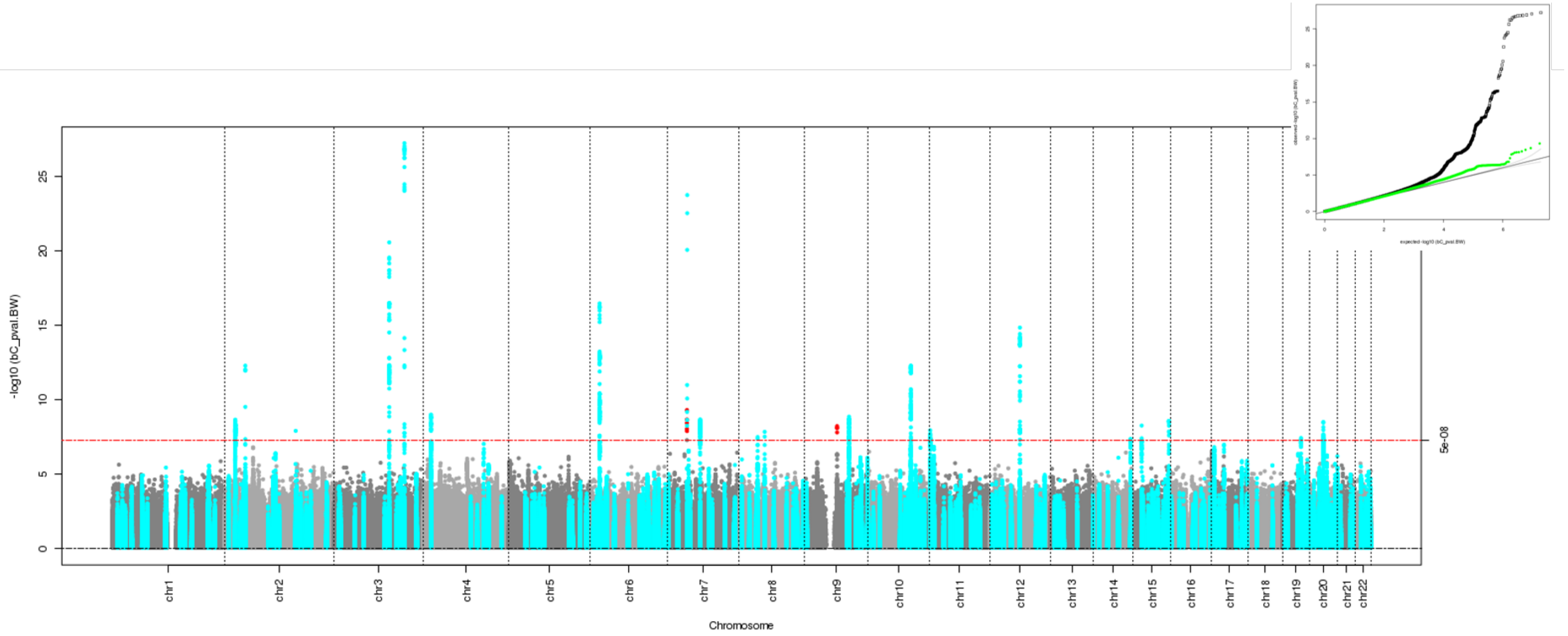

**Supplementary Figure 13: Manhattan plot and quantile-quantile (Q-Q) plot for the offspring specific effect estimated Genomic SEM[4] and sample overlap between the GWAS of own and offspring birth weight.** The two-sided association P-value, on the –log10 scale, obtained from Genomic SEM for each of the SNPs (y-axis) was plotted against the genomic position (NCBI Build 37; x-axis). Association signals that reached genome-wide significance ($P < 5x10^{-8}$) are shown in red if they are novel and turquoise if they have been previously reported[1]. In the Q-Q plots, the black dots represent observed P-values and the grey line represents expected P-values under the null distribution. The green dots represent observed P-values after excluding the previously identified signals[1].

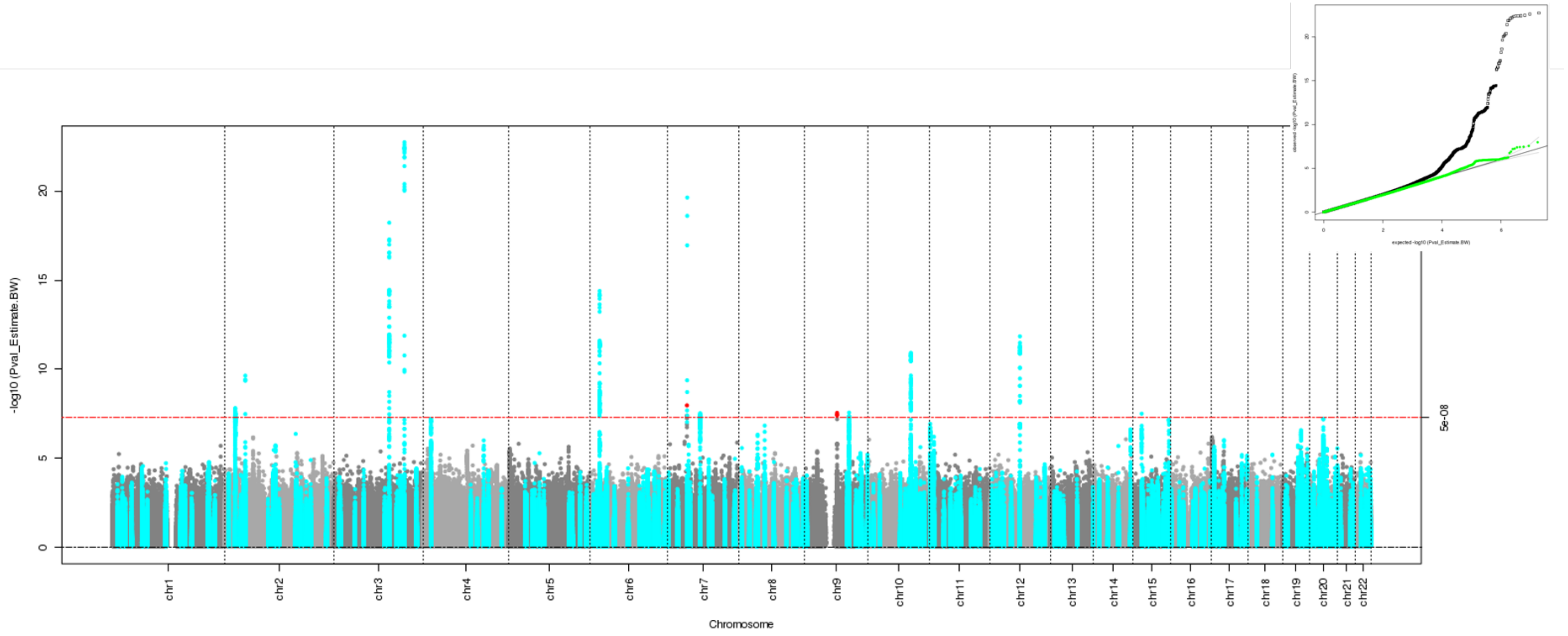

**Supplementary Figure 14: Manhattan plot and quantile-quantile (Q-Q) plot for the maternal specific effect estimated using the structural equation model (SEM) with summary statistics and no sample overlap between the GWAS of own and offspring birth weight.** The two-sided association P-value, on the –log10 scale, obtained from the SEM for each of the SNPs (y-axis) was plotted against the genomic position (NCBI Build 37; x-axis). Association signals that reached genome-wide significance ($P < 5x10^{-8}$) are shown in red if they are novel and turquoise if they have been previously reported[1]. In the Q-Q plots, the black dots represent observed P-values and the grey line represents expected P-values under the null distribution. The green dots represent observed P-values after excluding the previously identified signals[1].

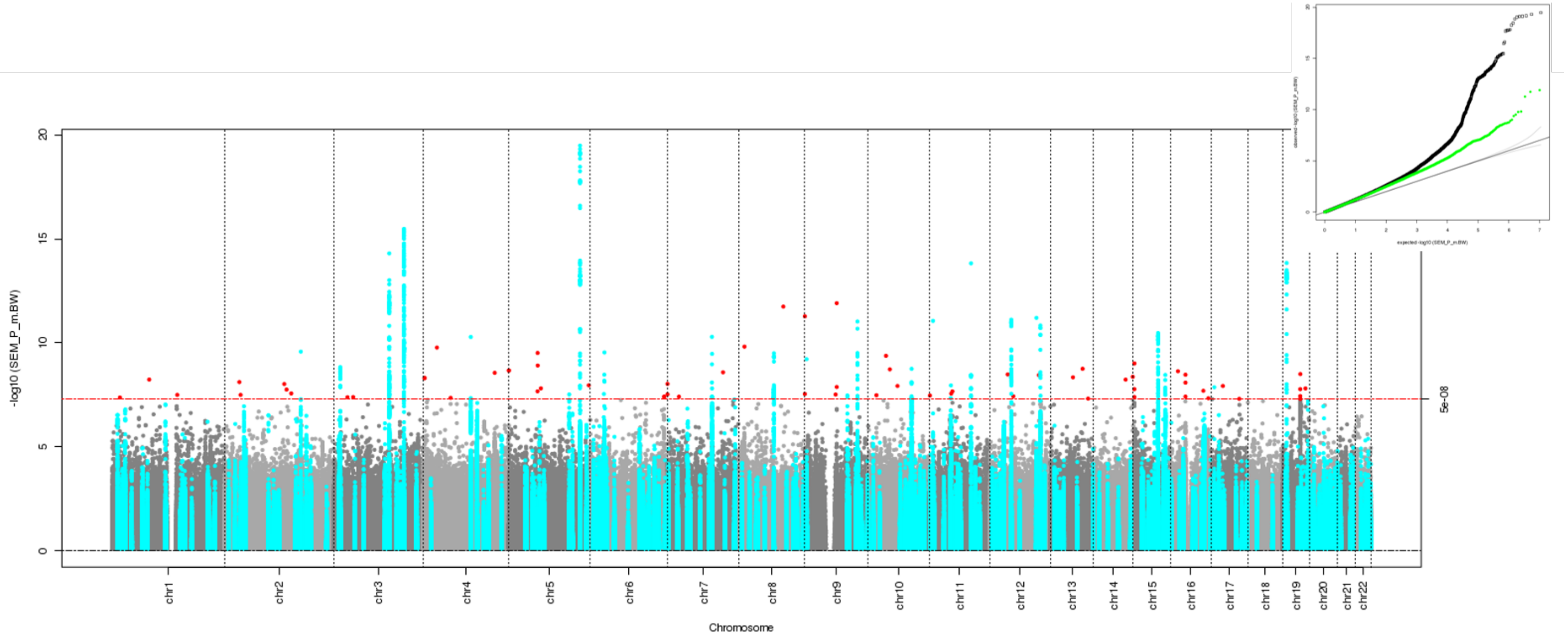

**Supplementary Figure 15: Manhattan plot and quantile-quantile (Q-Q) plot for the maternal specific effect estimated using the linear approximation of the structural equation model (SEM) and no sample overlap between the GWAS of own and offspring birth weight.** The two-sided association P-value, on the –log10 scale, obtained from the linear approximation of the SEM for each of the SNPs (y-axis) was plotted against the genomic position (NCBI Build 37; x-axis). Association signals that reached genome-wide significance ($P < 5x10^{-8}$) are shown in red if they are novel and turquoise if they have been previously reported[1]. In the Q-Q plots, the black dots represent observed P-values and the grey line represents expected P-values under the null distribution. The green dots represent observed P-values after excluding the previously identified signals[1].

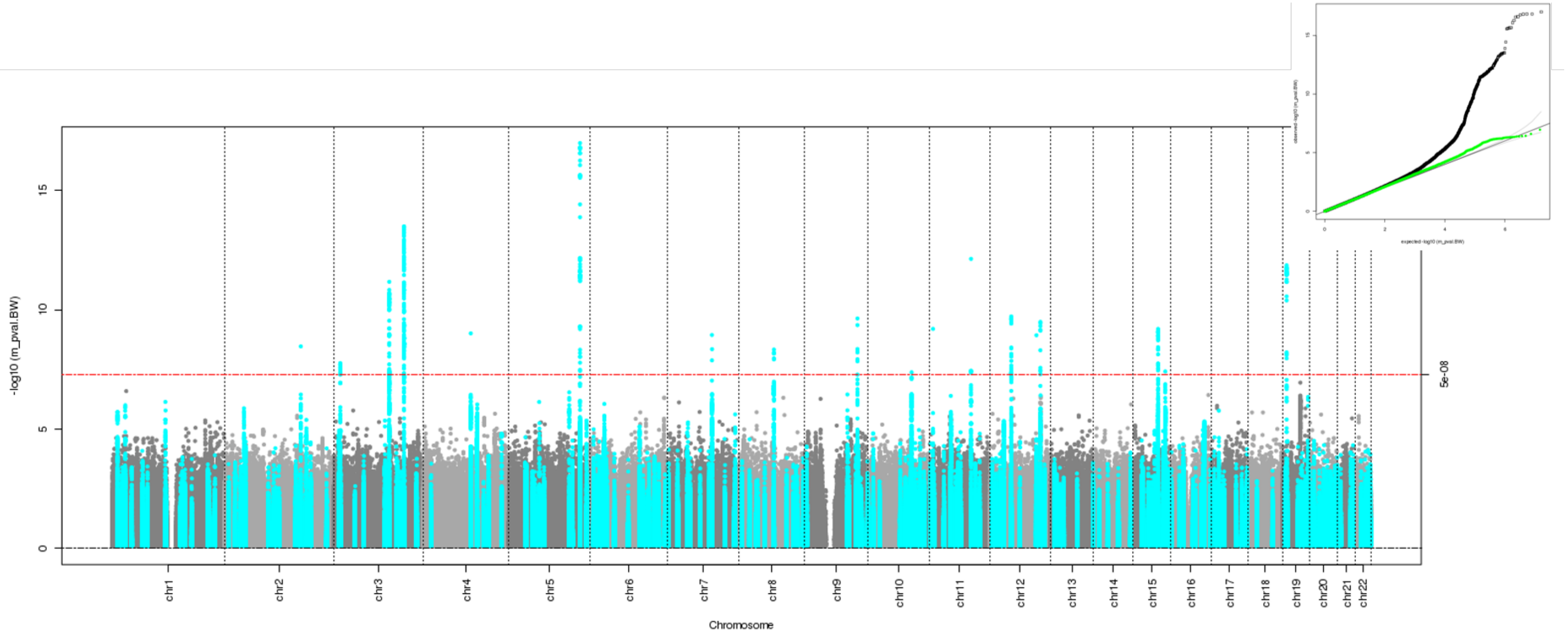

**Supplementary Figure 16: Manhattan plot and quantile-quantile (Q-Q) plot for the maternal specific effect estimated using MTAG[2] and no sample overlap between the GWAS of own and offspring birth weight.** The two-sided association P-value, on the –log10 scale, obtained from MTAG for each of the SNPs (y-axis) was plotted against the genomic position (NCBI Build 37; x-axis). Association signals that reached genome-wide significance ($P < 5x10^{-8}$) are shown in red if they are novel and turquoise if they have been previously reported[1]. In the Q-Q plots, the black dots represent observed P-values and the grey line represents expected P-values under the null distribution. The green dots represent observed P-values after excluding the previously identified signals[1].

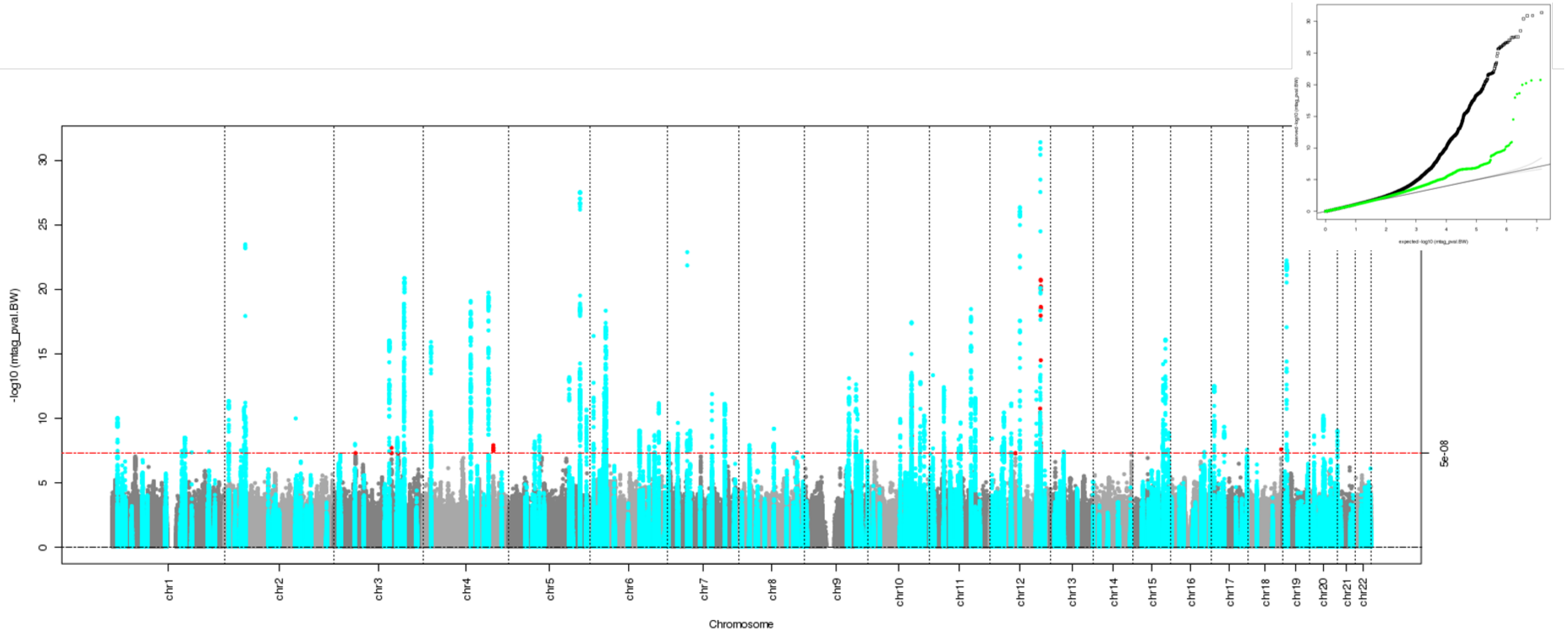

**Supplementary Figure 17: Manhattan plot and quantile-quantile (Q-Q) plot for the maternal specific effect estimated using mtCOJO[3] and no sample overlap between the GWAS of own and offspring birth weight.** The two-sided association P-value, on the –log10 scale, obtained from mtCOJO for each of the SNPs (y-axis) was plotted against the genomic position (NCBI Build 37; x-axis). Association signals that reached genome-wide significance ($P < 5x10^{-8}$) are shown in red if they are novel and turquoise if they have been previously reported[1]. In the Q-Q plots, the black dots represent observed P-values and the grey line represents expected P-values under the null distribution. The green dots represent observed P-values after excluding the previously identified signals[1].

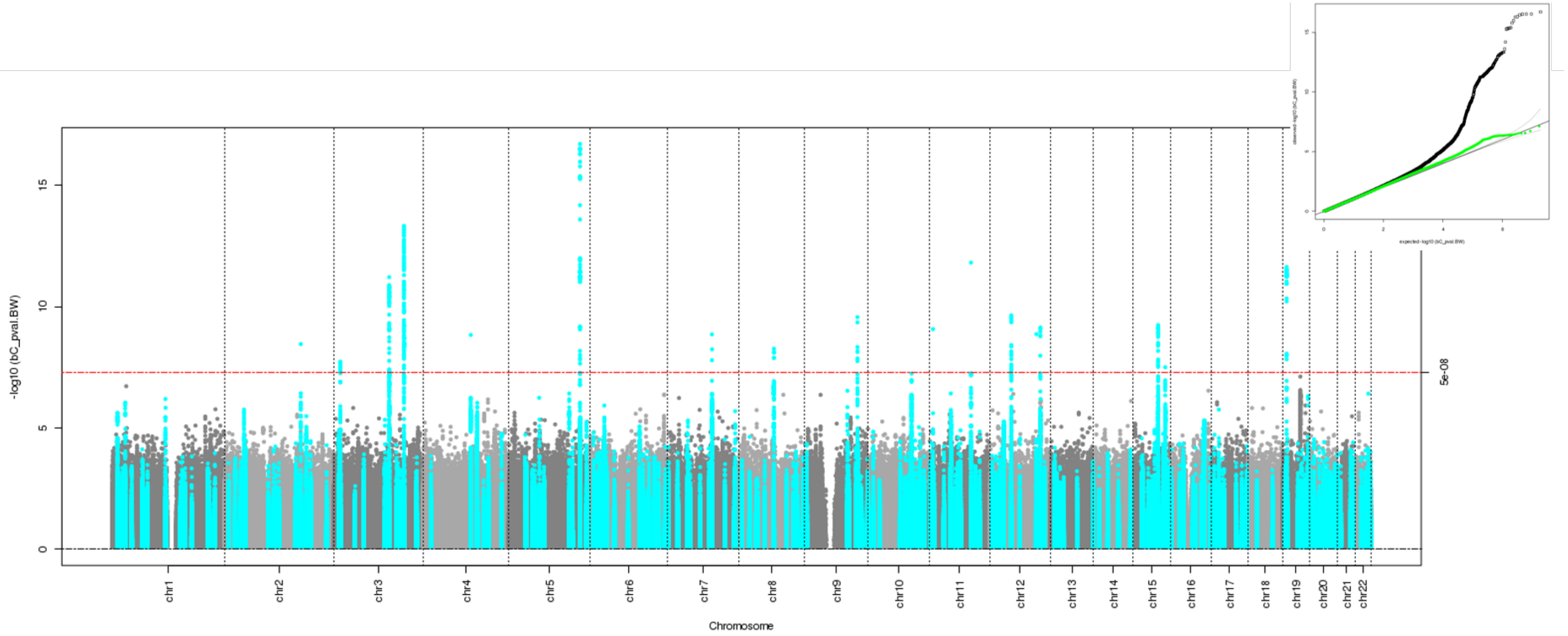

**Supplementary Figure 18: Manhattan plot and quantile-quantile (Q-Q) plot for the maternal specific effect estimated using Genomic SEM[4] and no sample overlap between the GWAS of own and offspring birth weight.** The two-sided association P-value, on the –log10 scale, obtained from Genomic SEM for each of the SNPs (y-axis) was plotted against the genomic position (NCBI Build 37; x-axis). Association signals that reached genome-wide significance ($P < 5x10^{-8}$) are shown in red if they are novel and turquoise if they have been previously reported[1]. In the Q-Q plots, the black dots represent observed P-values and the grey line represents expected P-values under the null distribution. The green dots represent observed P-values after excluding the previously identified signals[1].

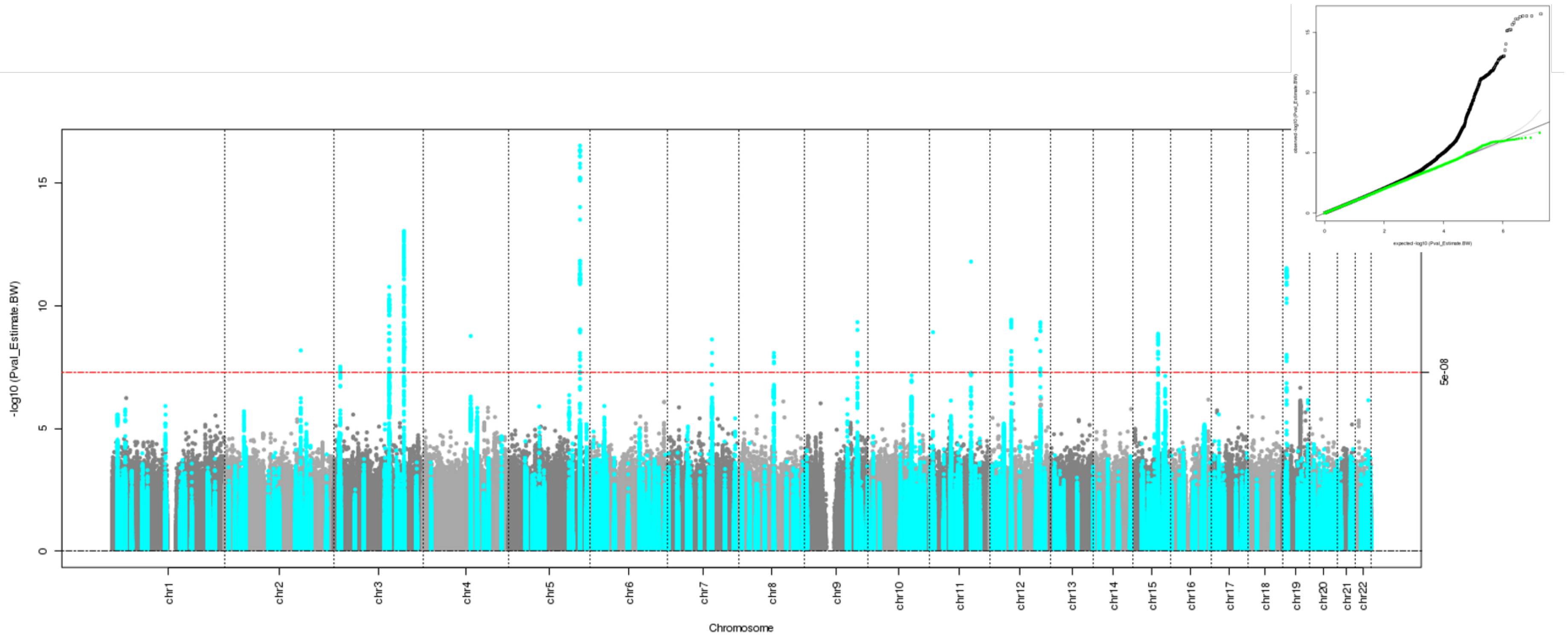

**Supplementary Figure 19: Manhattan plot and quantile-quantile (Q-Q) plot for the maternal specific effect estimated using the structural equation model (SEM) with summary statistics and sample overlap between the GWAS of own and offspring birth weight.** The two-sided association P-value, on the –log10 scale, obtained from the SEM for each of the SNPs (y-axis) was plotted against the genomic position (NCBI Build 37; x-axis). Association signals that reached genome-wide significance ($P < 5x10^{-8}$) are shown in red if they are novel and turquoise if they have been previously reported[1]. In the Q-Q plots, the black dots represent observed P-values and the grey line represents expected P-values under the null distribution. The green dots represent observed P-values after excluding the previously identified signals[1].

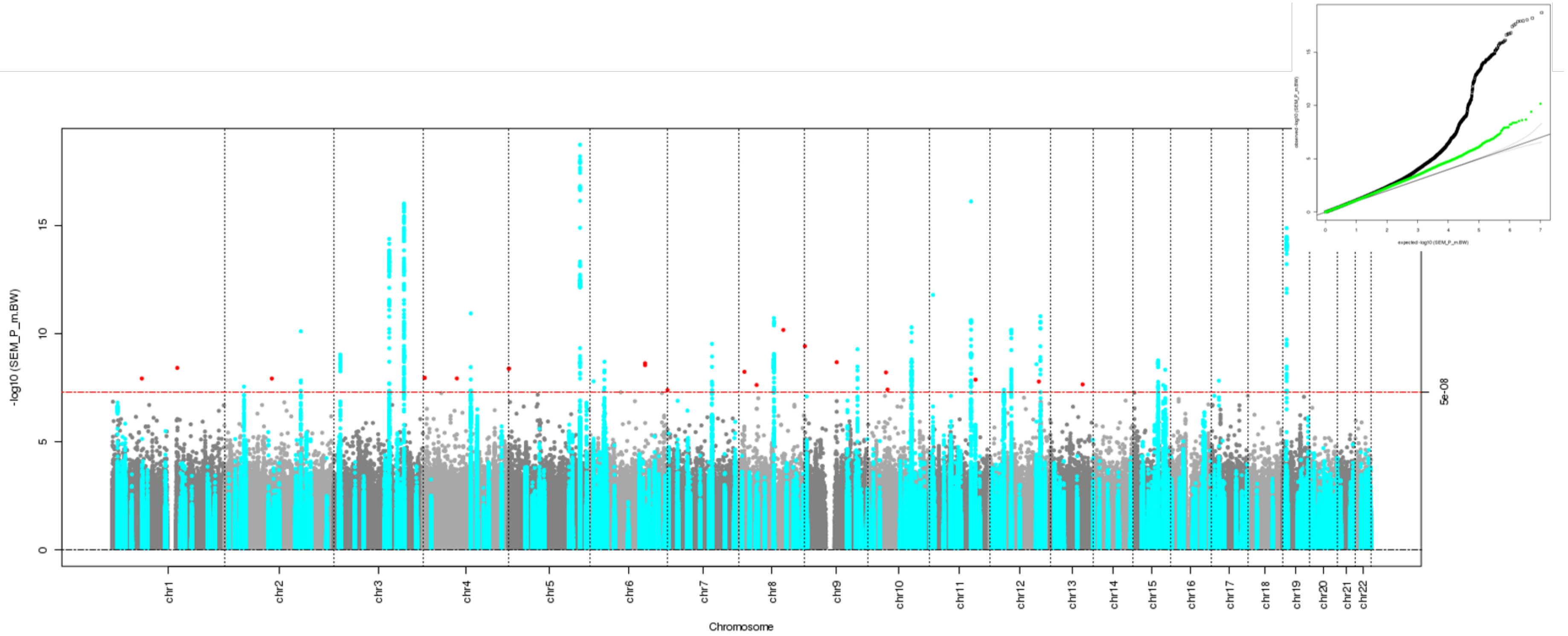

**Supplementary Figure 20: Manhattan plot and quantile-quantile (Q-Q) plot for the maternal specific effect estimated using the linear approximation of the structural equation model (SEM) and sample overlap between the GWAS of own and offspring birth weight.** The two-sided association P-value, on the –log10 scale, obtained from the linear approximation of the SEM for each of the SNPs (y-axis) was plotted against the genomic position (NCBI Build 37; x-axis). Association signals that reached genome-wide significance ($P < 5x10^{-8}$) are shown in red if they are novel and turquoise if they have been previously reported[1]. In the Q-Q plots, the black dots represent observed P-values and the grey line represents expected P-values under the null distribution. The green dots represent observed P-values after excluding the previously identified signals[1].

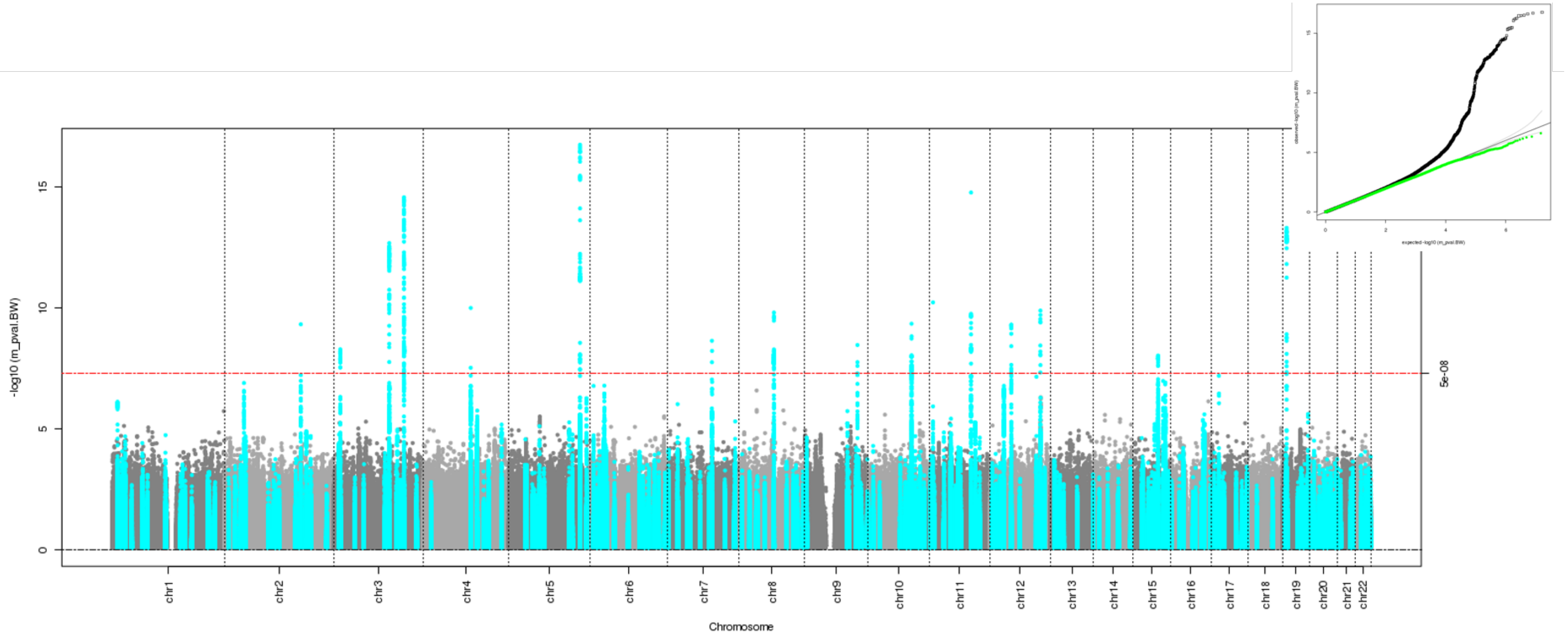

**Supplementary Figure 21: Manhattan plot and quantile-quantile (Q-Q) plot for the maternal specific effect estimated using MTAG[2] and sample overlap between the GWAS of own and offspring birth weight.** The two-sided association P-value, on the –log10 scale, obtained from MTAG for each of the SNPs (y-axis) was plotted against the genomic position (NCBI Build 37; x-axis). Association signals that reached genome-wide significance ($P < 5x10^{-8}$) are shown in red if they are novel and turquoise if they have been previously reported[1]. In the Q-Q plots, the black dots represent observed P-values and the grey line represents expected P-values under the null distribution. The green dots represent observed P-values after excluding the previously identified signals[1].

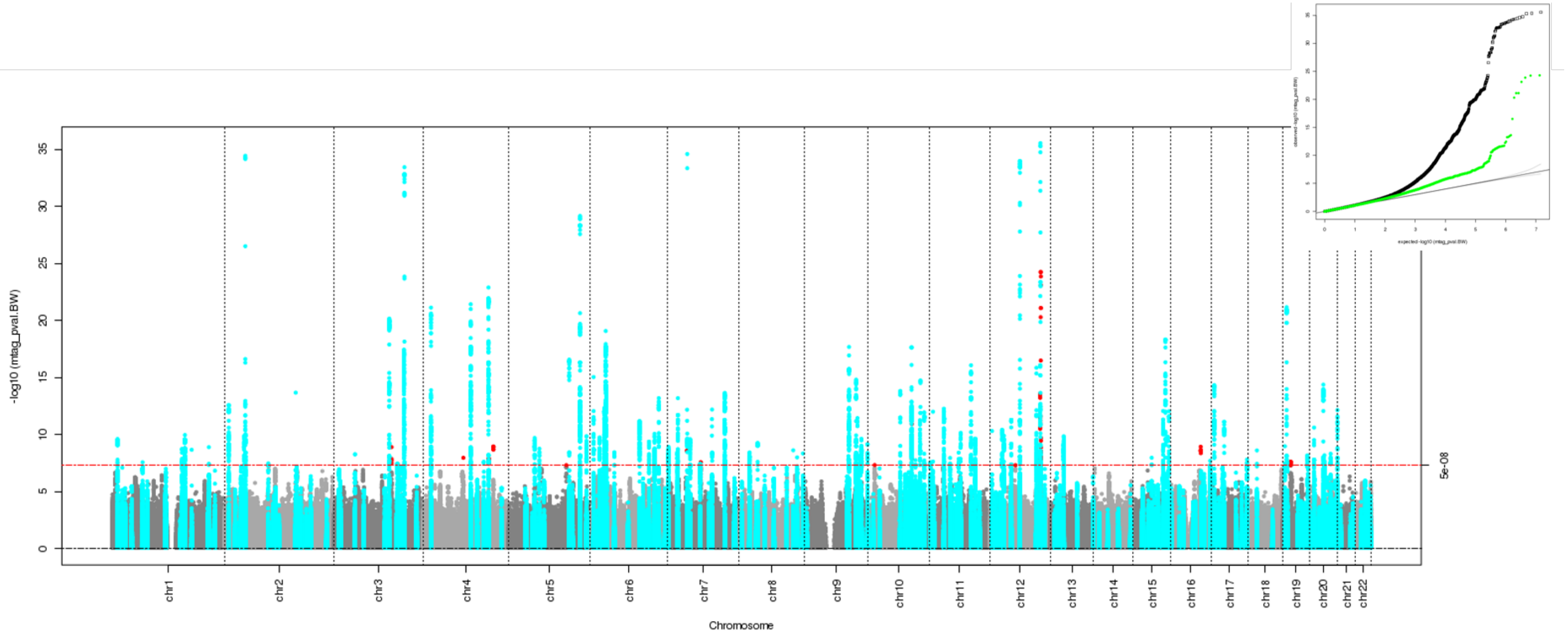

**Supplementary Figure 22: Manhattan plot and quantile-quantile (Q-Q) plot for the maternal specific effect estimated mtCOJO[3] and sample overlap between the GWAS of own and offspring birth weight.** The two-sided association P-value, on the –log10 scale, obtained from mtCOJO for each of the SNPs (y-axis) was plotted against the genomic position (NCBI Build 37; x-axis). Association signals that reached genome-wide significance ($P < 5x10^{-8}$) are shown in red if they are novel and turquoise if they have been previously reported[1]. In the Q-Q plots, the black dots represent observed P-values and the grey line represents expected P-values under the null distribution. The green dots represent observed P-values after excluding the previously identified signals[1].

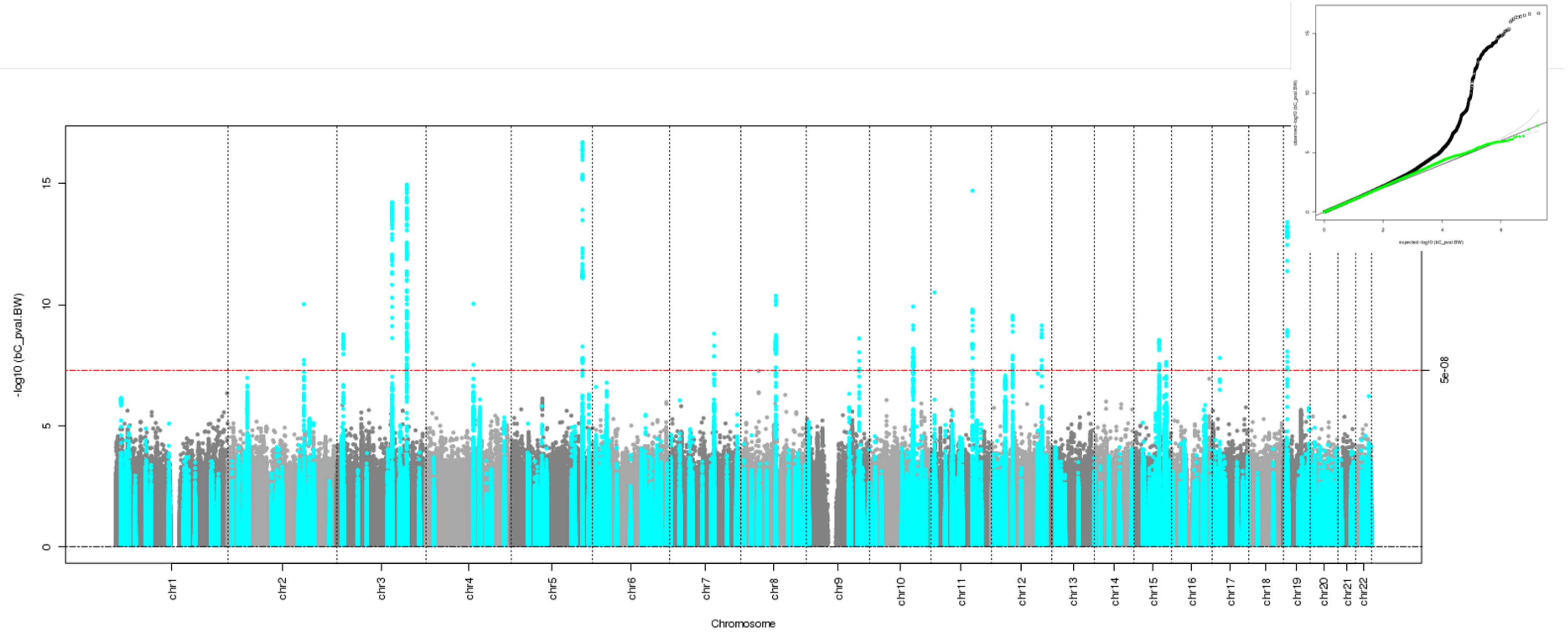

**Supplementary Figure 23: Manhattan plot and quantile-quantile (Q-Q) plot for the maternal specific effect estimated using Genomic SEM[4] and sample overlap between the GWAS of own and offspring birth weight.** The two-sided association P-value, on the –log10 scale, obtained from Genomic SEM for each of the SNPs (y-axis) was plotted against the genomic position (NCBI Build 37; x-axis). Association signals that reached genome-wide significance ($P < 5x10^{-8}$) are shown in red if they are novel and turquoise if they have been previously reported[1]. In the Q-Q plots, the black dots represent observed P-values and the grey line represents expected P-values under the null distribution. The green dots represent observed P-values after excluding the previously identified signals[1].

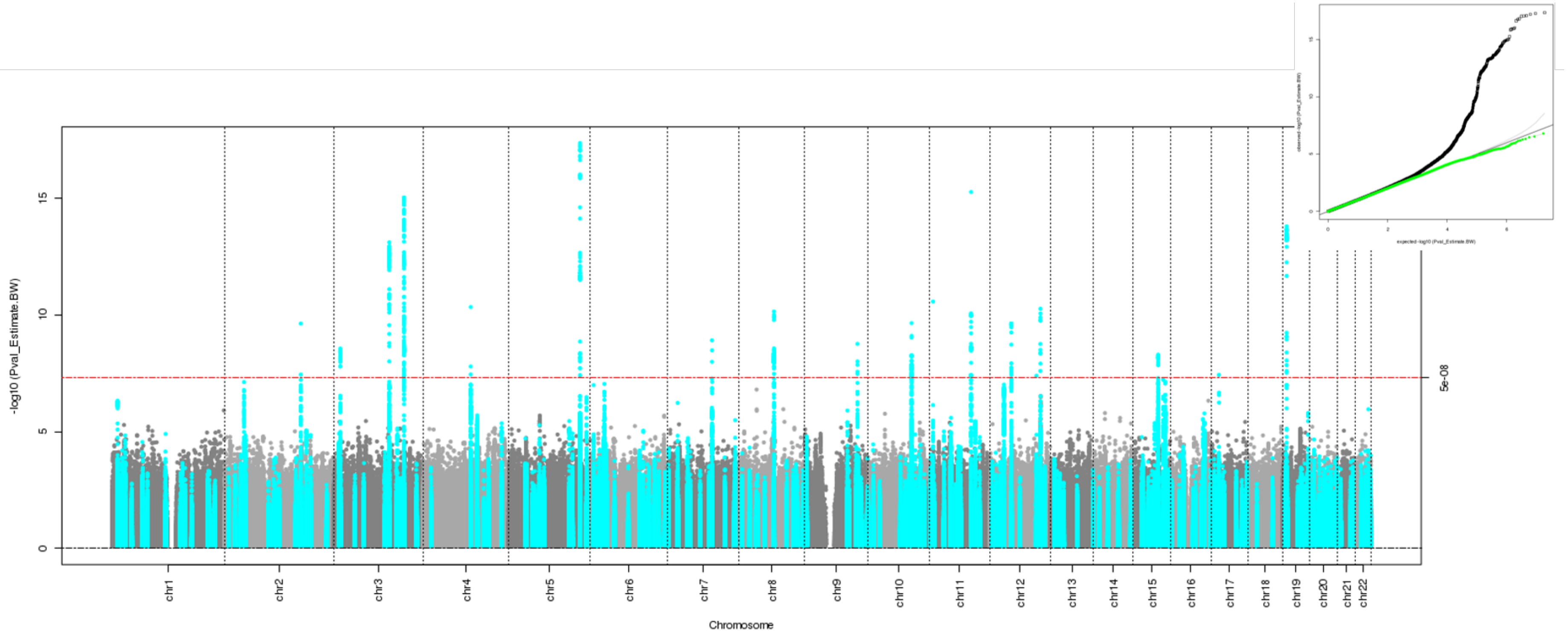

**Supplementary Figure 24: Quantile-quantile (Q-Q) plot for the offspring and maternal specific effects estimated using mtCOJO using simulated data, both with and without sample overlap between the GWAS of own and offspring birth weight.**

| | Offspring specific effect | Maternal specific effect |
|---|---|---|
| No sample overlap | 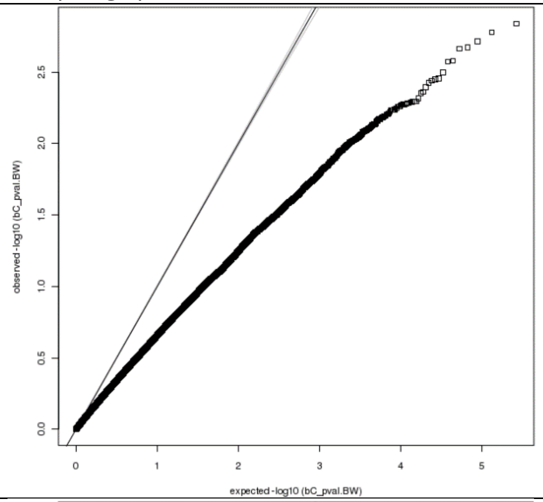 | 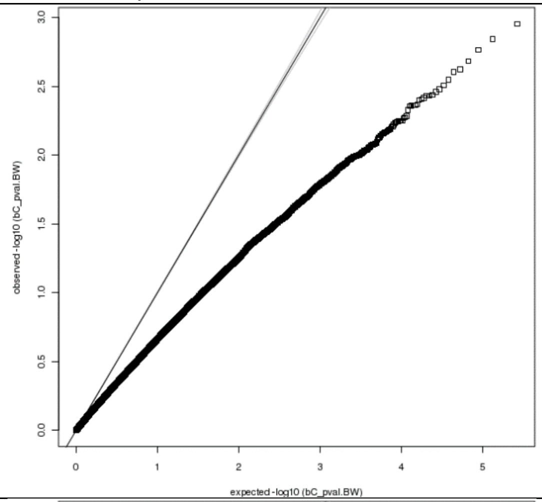 |
| Sample overlap | 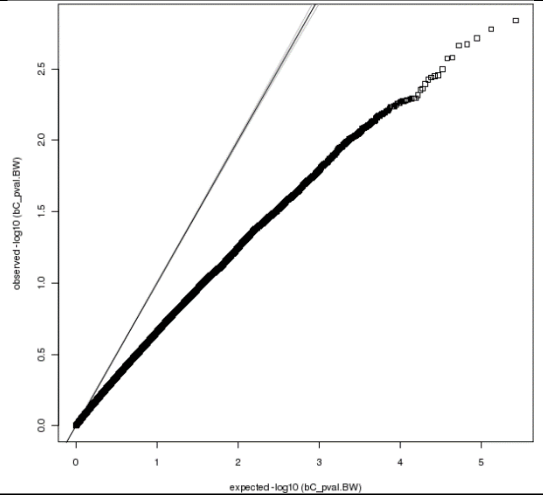 | 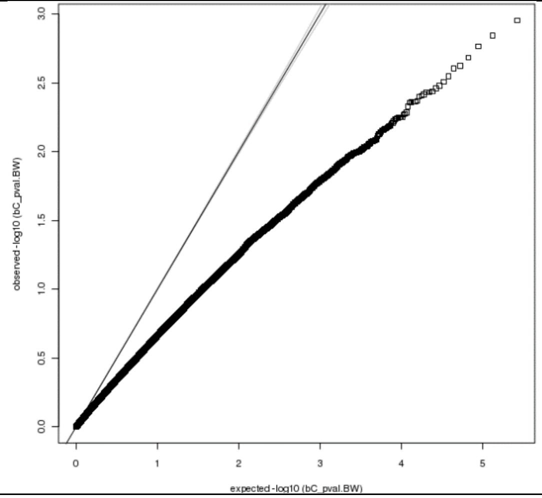 |

**Supplementary Figure 25: Comparison of the offspring effect size estimates and the standard errors using the linear approximation of the structural equation model (SEM; x-axis) and mtCOJO (y-axis) for the genetic variants on chromosome 22 using simulated birth weight data.**

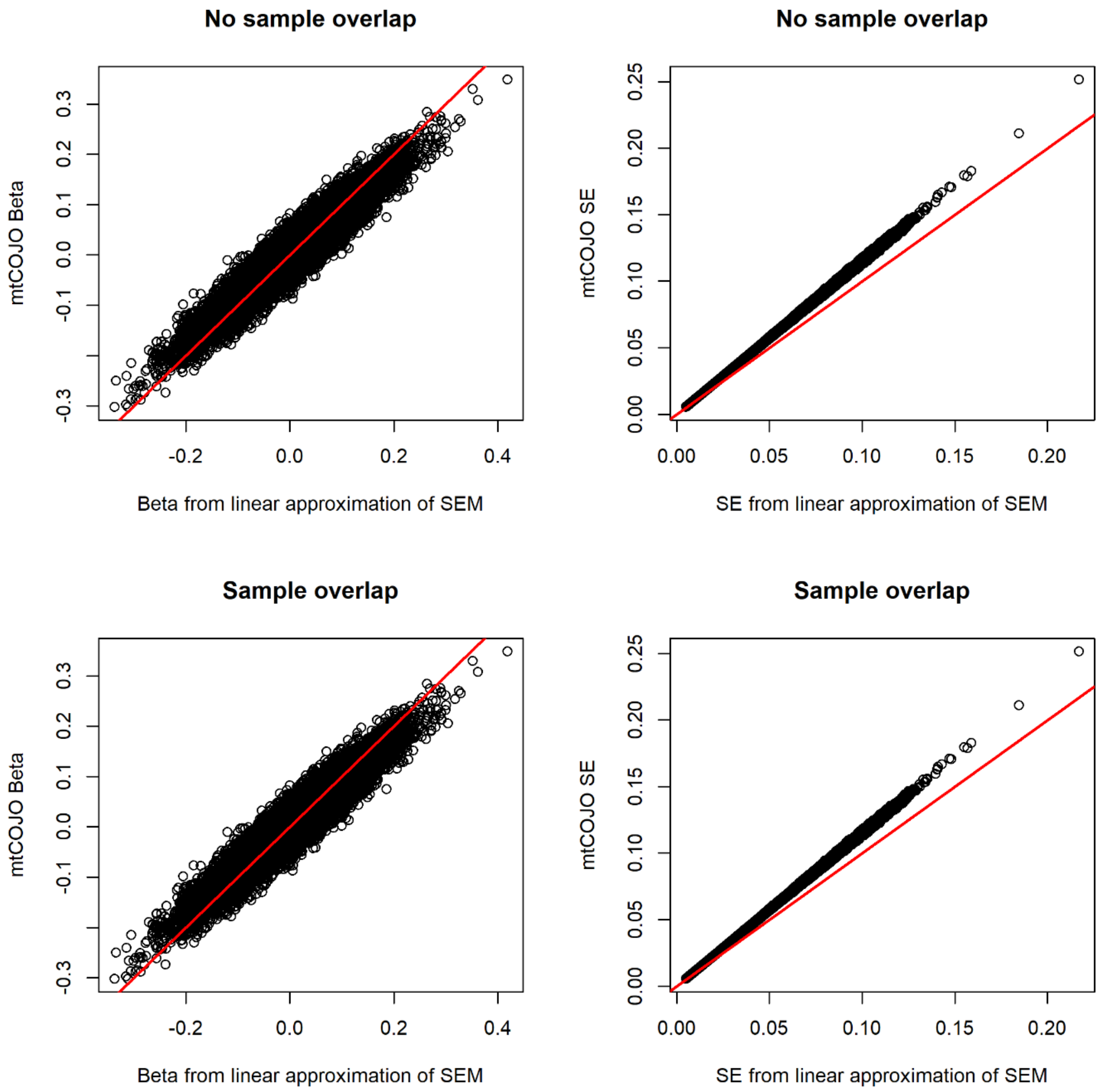

**Supplementary Figure 26: Comparison of the maternal effect size estimates and the standard errors using the linear approximation of the structural equation model (SEM; x-axis) and mtCOJO (y-axis) for the genetic variants on chromosome 22 using simulated birth weight data.**

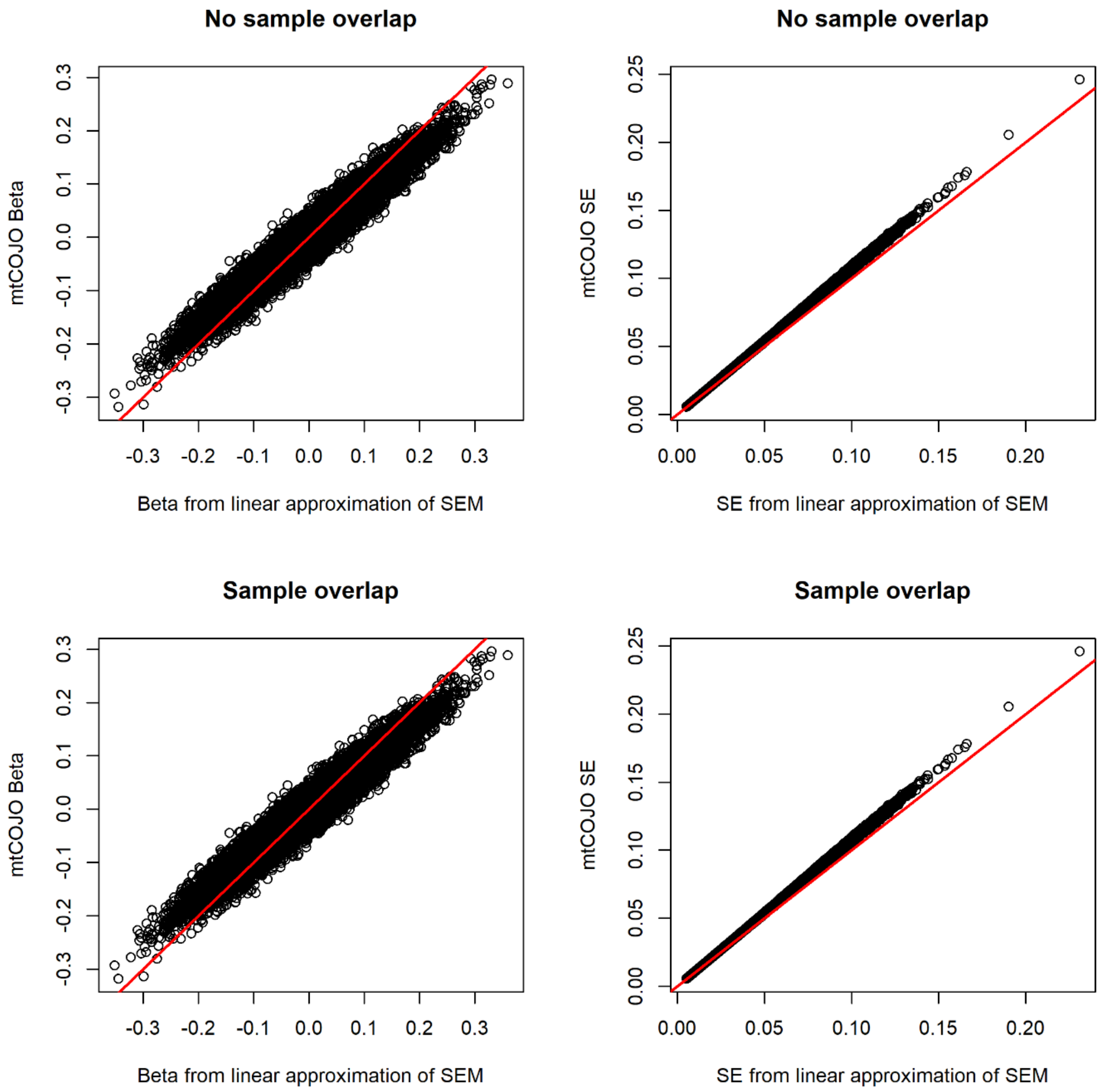

**Supplementary Figure 27: Manhattan plot and quantile-quantile (Q-Q) plot for the fertility GWAS estimating the genetic effects on number of children mothered and fathered and the number of siblings estimated using BOLT-LMM.** The two-sided association P-value, on the –log10 scale, obtained from BOLT-LMM for each of the SNPs (y-axis) was plotted against the genomic position (NCBI Build 37; x-axis). Association signals that reached genome-wide significance ($P < 5x10^{-8}$) are shown in red. In the Q-Q plots, the black dots represent observed P-values and the grey line represents expected P-values under the null distribution.

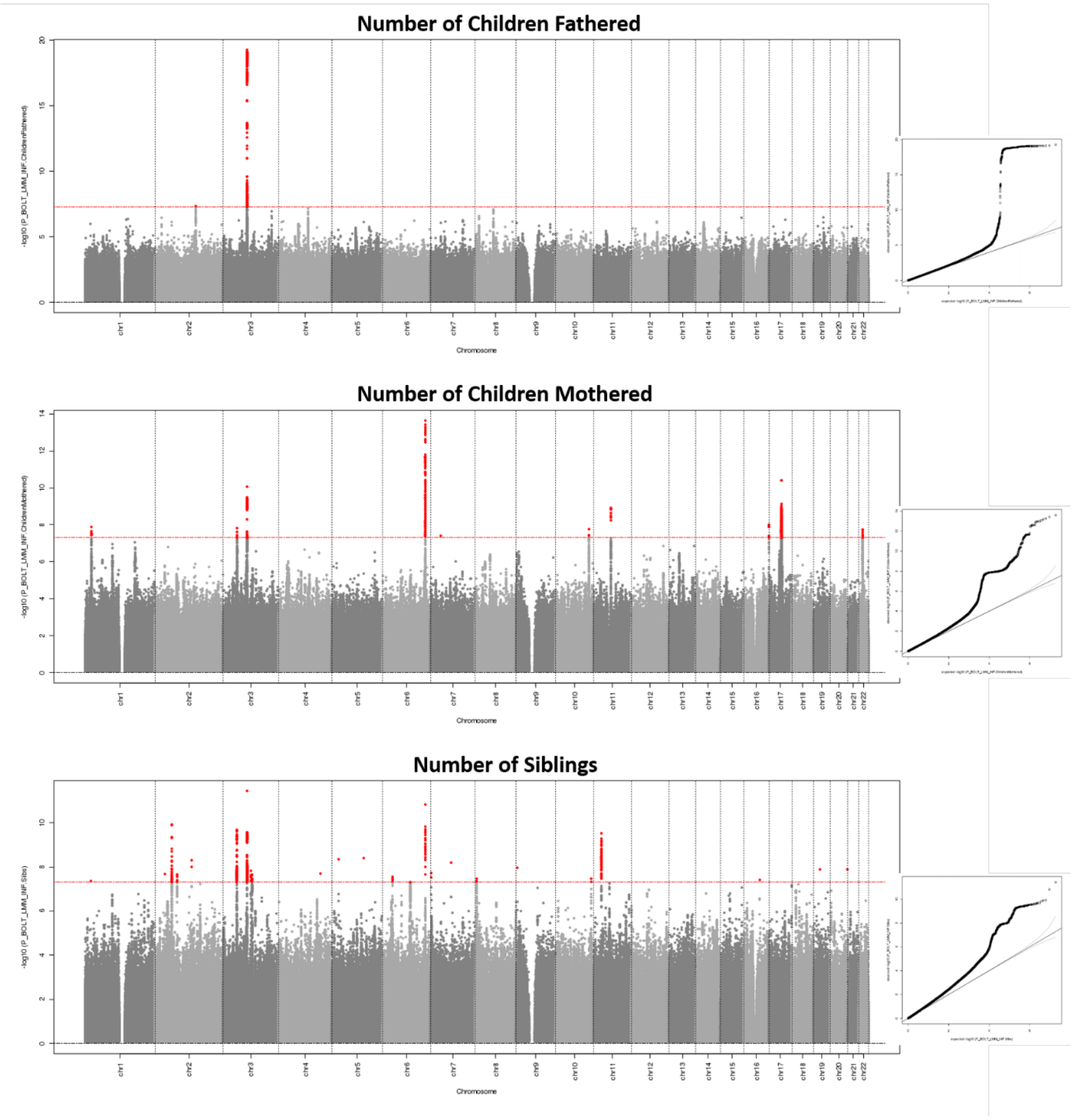

**Supplementary Figure 28: Manhattan plot and quantile-quantile (Q-Q) plot for the fertility GWAS estimating maternal and offspring specific genetic effects using Genomic SEM.** The two-sided association P-value, on the –log10 scale, obtained from Genomic SEM for each of the SNPs (y-axis) was plotted against the genomic position (NCBI Build 37; x-axis). Association signals that reached genome-wide significance ($P < 5x10^{-8}$) are shown in red. In the Q-Q plots, the black dots represent observed P-values and the grey line represents expected P-values under the null distribution.

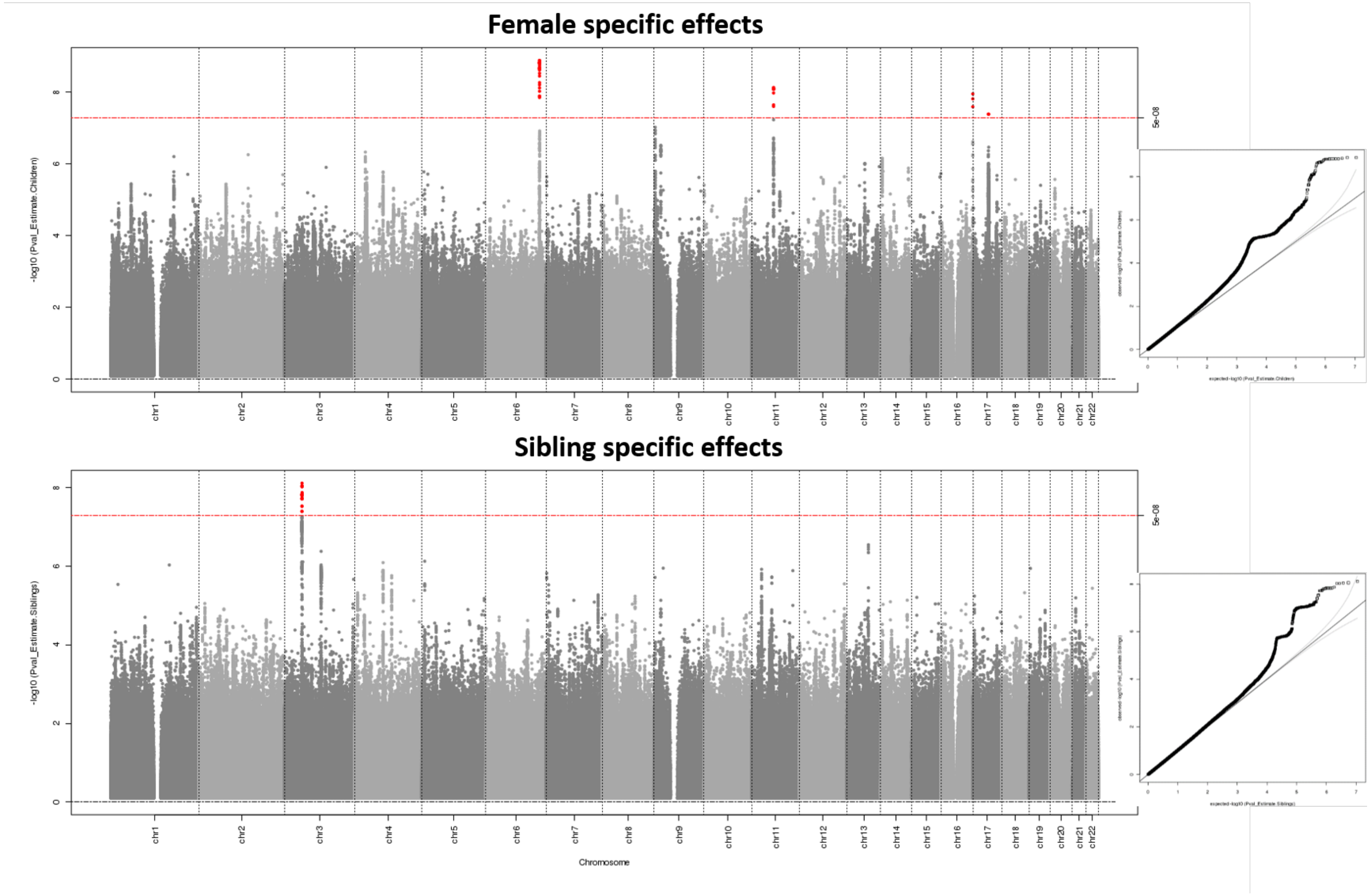

**Supplementary Figure 29: Genetic correlation between male and female fertility and sibling specific effects and traits related to development, reproduction, behaviour, neuropsychiatric disorders and anthropometry.** Genetic correlations were calculated using LD score regression[5], conducted in LD Hub[6]. The traits were chosen based on those used in Barban et al[7]. The point indicates the genetic correlation and the bars indicate the 95% confidence interval; the size of the point is proportional to 1/(standard error)$^2$. The mark "*" indicates that the estimate of genetic correlation is statistically significant after controlling for multiple testing (P<0.05/22=0.002).

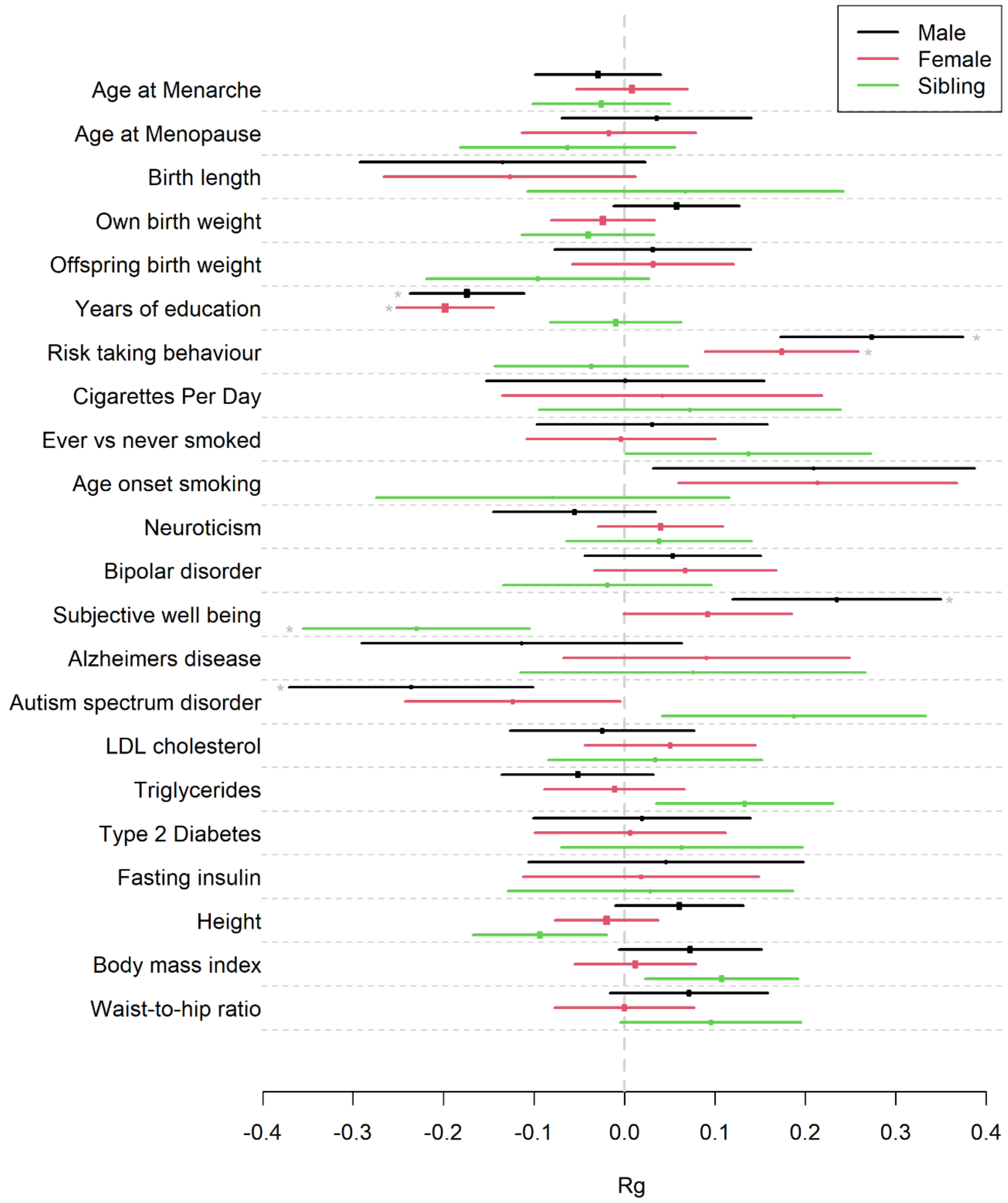
